# Endogenous activation of multiple stress-response pathways during meiosis promotes gamete fitness

**DOI:** 10.64898/2026.11.18.767631

**Authors:** Alena L. Bishop, Silvan Spiri, Defne Yalçın, Gloria A. Brar, Elçin Ünal

## Abstract

Gametogenesis, the process of meiosis coupled to gamete formation, transforms diploid progenitor cells into haploid gametes through extensive cellular remodeling and rewiring of gene expression. Several stress-responsive transcription factors (TFs) are upregulated during gametogenesis, but whether the cellular mechanisms that handle environmental stress also support the success of this developmental process is largely unexplored. Here, we show that four stress-responsive TFs, Hsf1, Skn7, and the paralogs Msn2 and Msn4, are endogenously active during budding yeast gametogenesis in the absence of external stress and drive distinct gene expression programs. Simultaneous meiotic depletion of Msn2, Msn4, and Skn7 compromise the fitness of the resulting gametes. We identified just two TF-dependent targets, the antioxidant genes *TSA1* and *AHP1*, as important for gamete fitness, directly connecting TF activity to the protection of meiotic progeny. Our results demonstrate that the activation of canonical stress-response regulators is built into the meiotic program and suggest that such pathways can be repurposed during development to safeguard cellular health.

## Introduction

Gametogenesis is a conserved developmental program consisting of meiosis coupled with gamete construction that transforms diploid progenitor cells into haploid gametes capable of giving rise to the next generation. Production of functional gametes requires the coordination of chromosome segregation with extensive changes in gene expression, metabolism, and cellular architecture, including large-scale membrane remodeling and a requirement for aerobic respiration^1–14^. Such extensive remodeling can itself create physiological challenges that must be managed while progenitor cells differentiate into gametes. How cells maintain homeostasis throughout differentiation, and whether pathways canonically associated with environmental stress contribute to this process, remain poorly understood.

Across eukaryotes, stress-responsive transcriptional pathways preserve cellular homeostasis in the face of environmental and physiological perturbations. These pathways have traditionally been studied following acute challenges such as heat shock, oxidative stress, nutrient limitation, or protein misfolding, and consequently are often viewed primarily as mechanisms that enable cells to survive adverse conditions^15–20^. However, increasing evidence indicates that canonical stress-response pathways also perform physiological functions during development and differentiation. For example, HSF1 drives a transcriptional program required for *C. elegans* larval development that is distinct from the canonical heat-shock response^21^, and the unfolded protein response is physiologically activated during mammalian plasma-cell differentiation and contributes to the development of antibody-secreting cells^22,23^. These observations suggest that stress-response pathways can be incorporated into normal developmental programs, but how multiple such pathways operate within a developmental program and how they influence the fitness of the resulting differentiated cells remain largely unexplored.

Gametogenesis in budding yeast provides a powerful setting in which to address these questions. Rather than engaging a single homeostatic pathway, gametogenesis is accompanied by the endogenous activation of multiple stress-responsive transcription factors (TFs) in the absence of overt environmental stress^6,24^. These include Hsf1, a conserved regulator of proteostasis and the heat-shock response; Skn7, which contributes to the oxidative-stress response; and the paralogous TFs Msn2 and Msn4, which respond to diverse environmental perturbations^15,19^. We previously found that another canonical stress-response pathway, the unfolded protein response, is physiologically activated to prepare cells for entry into gametogenesis^25^. Thus, gametogenesis offers an opportunity to determine how distinct stress-response programs are developmentally utilized within the same cellular differentiation program, whether they serve overlapping or specialized functions, and how their activity ultimately affects the health of the resulting gametes.

Here, we investigated the endogenous functions of Hsf1, Skn7, Msn2, and Msn4 during budding yeast gametogenesis. By conducting high-resolution time course experiments, we found that these TFs execute distinct transcriptional programs during gametogenesis. Further, we show that meiotic depletion of a subset of these factors compromises the fitness of the resulting gametes. We identified two TF-dependent antioxidant genes, *TSA1* and *AHP1*, that together are required for gamete fitness, linking the endogenous activity of stress-responsive TFs to the protection of meiotic progeny. Together, our findings establish that canonical stress-response regulators are developmentally used during gametogenesis to promote gamete health. More broadly, they reveal how transcriptional pathways traditionally associated with responding to environmental insults can be integrated into developmental programs to preserve cellular fitness through cell-state transitions and ultimately safeguard the ensuing progeny.

## Results

### Stress-responsive TFs are dynamically expressed during gametogenesis and can be acutely depleted

To investigate the functions of Hsf1, Skn7, Msn2, and Msn4 during gametogenesis, we used the AID2 system^26^ to acutely deplete each TF after cells were induced to undergo gametogenesis. We tagged each TF at its endogenous locus with a 3V5 epitope and the IAA17 auxin-inducible degron (hereafter AID), enabling detection and conditional degradation of the tagged proteins. To restrict depletion to gametogenesis, we expressed OsTIR1^F74G^ from the copper-inducible *P_CUP1_* promoter and induced degradation by addition of CuSO_4_ and the auxin analog 5-Ph-IAA (Fig. 1A). Control strains lacking OsTIR1^F74G^ retained TF expression following CuSO_4_ and 5-Ph-IAA treatment (Fig. 1A). Because Msn2 and Msn4 are paralogous regulators of the general stress response, we tagged both proteins in the same strain to enable their simultaneous depletion.

**Fig. 1.**
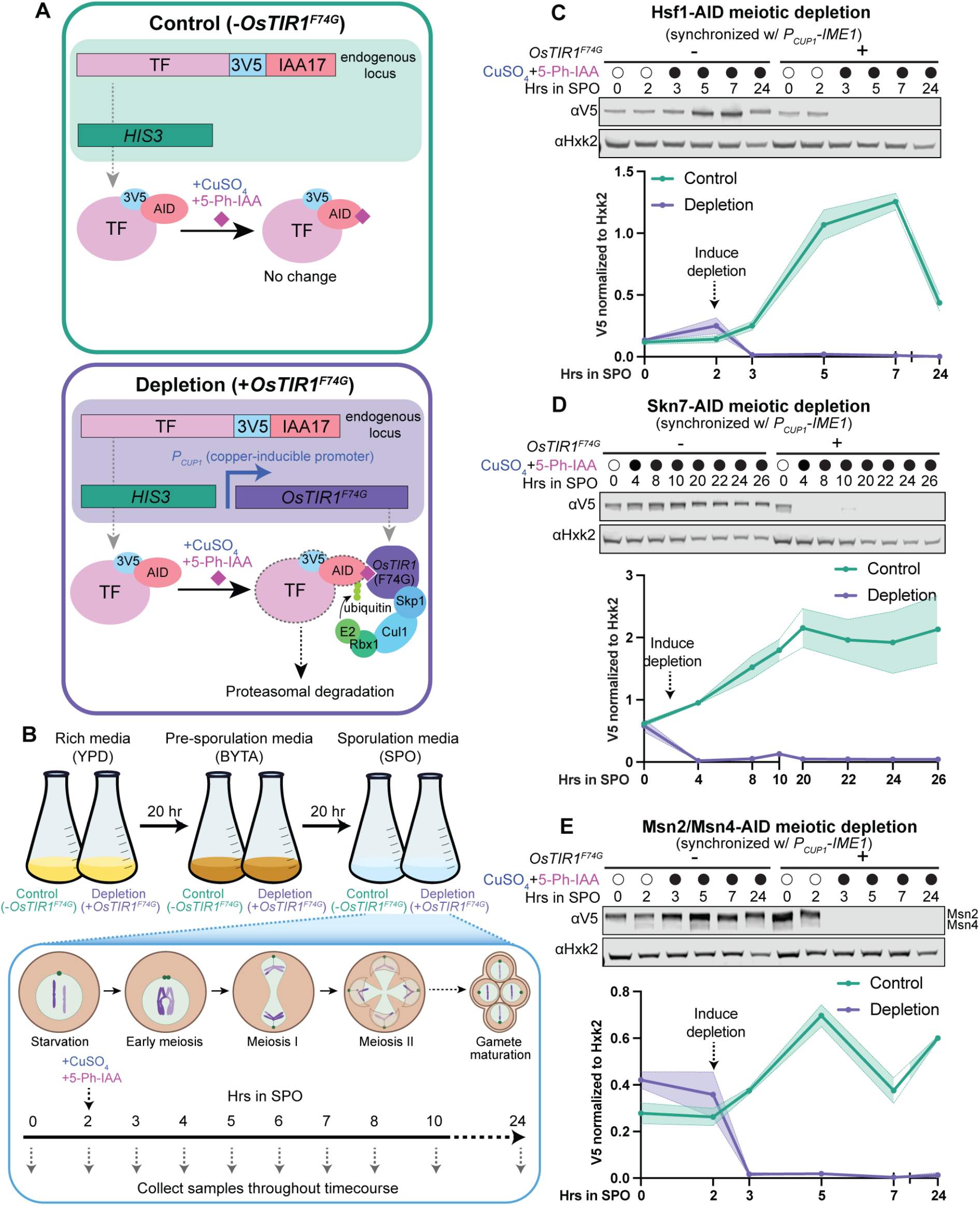
Stress-responsive TFs are dynamically expressed during gametogenesis and can be acutely depleted. **A** Schematics of control (top) and depletion (bottom) conditions using AID2 system. Genotypes are shown in shaded regions. In both strains, the TF is tagged at the endogenous locus with a 3V5 epitope for immunoblotting and IAA17 degron for depletion (hereafter AID). Control and depletion strains are prototrophy matched, but only the depletion strains contain *OsTIR1^F74G^* under the copper-inducible *P_CUP1_* promoter at the *HIS3* locus. Upon addition of CuSO_4_ and the auxin analog 5-Ph-IAA, *OsTIR1^F74G^* is expressed and the tagged protein is ubiquitinated, targeting it for proteasomal degradation. **B** Schematic of experimental set up used to induce cells to enter gametogenesis in control and TF depletion conditions (top). Diploid cells are grown overnight in rich media (YPD), back diluted the next day in pre-sporulation media (BYTA), then transferred to nutritionally poor sporulation media (SPO) the following day to induce gametogenesis. Schematic of meiotic stages, with approximate timepoints at which they occur below (bottom). TF depletion is induced by addition of 50 µM CuSO_4_ and 100 µM 5-Ph-IAA at 2 hr in SPO, and protein samples are collected throughout the time course. **C-E** Western blots showing Hsf1-AID (C), Skn7-AID (D), and Msn2/Msn4-AID (E) protein levels throughout gametogenesis in control (-*OsTIR1^F74G^*) and depletion (+*OsTIR1^F74G^*) conditions. Msn4-AID is the faint, slightly smaller band below the brighter Msn2-AID band (E). Strains have the *P_CUP1_-IME1* allele, leading to synchronous meiotic entry upon addition of CuSO_4_. Hxk2 serves as a loading control. Quantification below each blot, shown is the average of 2 replicates, with shaded region representing the range.

We synchronized entry into gametogenesis, also known as sporulation in yeast, using the *P*_CUP1_-*IME1* system, which allows cells to be released into gametogenesis by addition of CuSO_4_ after 2 hr in sporulation medium (“SPO”, a nutritionally poor media that is permissive to meiotic entry)^27,28^. CuSO_4_ and 5-Ph-IAA were added at the time of meiotic induction, enabling depletion of each TF specifically during gametogenesis (Fig. 1B). Immunoblotting across a 24-hr meiotic time course revealed distinct temporal patterns of TF expression, consistent with our previous data for untagged proteins^6,24^. Hsf1 abundance peaked approximately 5-7 hr after transfer to SPO, whereas Skn7 levels peaked at 20 h, despite its sustained expression throughout meiosis (Fig. 1C,D). Msn2 and Msn4 reached their highest levels approximately 5 hr after transfer to SPO (Fig. 1E). In contrast, induction of the AID2 system resulted in rapid depletion of each TF to nearly undetectable levels that persisted throughout the meiotic time course (Fig. 1C-E). Thus, Hsf1, Skn7, Msn2, and Msn4 exhibit distinct expression dynamics during gametogenesis and can be acutely depleted following meiotic induction.

Because entry into gametogenesis is triggered by nutrient limitation, we next asked whether these expression dynamics reflect the meiotic program itself or simply the starvation conditions used to induce it. We therefore compared TF abundance in wild type (WT) *IME1* cells undergoing gametogenesis with that in *ime1*Δ cells exposed to the same sporulation conditions but unable to enter the meiotic program. In WT *IME1* cells, all four TFs displayed the dynamic expression patterns observed in our synchronized meiotic time courses. By contrast, TF levels were substantially less dynamic in *ime1*Δ cells (Fig. S1A-C). Hsf1 and Skn7 remained at relatively low and constant levels throughout starvation, whereas Msn2 and Msn4 increased modestly at later time points but lacked the pronounced dynamics observed in meiotic cells. The residual increase in Msn2 and Msn4 during starvation is consistent with their established roles in the general stress response. Together, these results indicate that the dynamic regulation of these stress-responsive TFs is coupled to the meiotic program rather than being solely a consequence of nutrient deprivation.

### Stress-responsive TFs activate temporally distinct transcriptional programs during gametogenesis

Having established a system to acutely deplete Hsf1, Skn7, and Msn2/Msn4 during gametogenesis, we next sought to define the meiotic transcriptional programs that they drive. Although these TFs and their targets have been extensively characterized during mitotic growth and in response to exogenous stress^19,29–40^, the genes they regulate during their endogenous activation in gametogenesis remain unknown. We therefore performed RNA-seq across meiotic time courses in control and TF-depleted cells, using synchronized meiotic entry to resolve TF-dependent changes in gene expression over time.

For Hsf1, we profiled 12 time points spanning 0-24 hr in SPO, with increased temporal resolution around 5 hr, when Hsf1 protein abundance begins to peak. Global gene expression patterns were largely preserved following Hsf1 depletion, with strong agreement between control and depletion conditions throughout most of gametogenesis (Pearson’s *r* ≥ 0.97) (Fig. 2A, S2A). The transcriptomes diverged most between 4 and 7 hr, and differential expression analysis of the 5, 5.5, and 6 hr time points identified 86 Hsf1-dependent genes (Fig. 3A). These included known HSf1 targets *SSA2*, encoding Hsp70^41^; *YDJ1* and *SIS1*, encoding Hsp40 co-chaperones^42,43^; and *HSP78*, encoding a mitochondrial chaperone^44^ (Fig. 3B). These genes were highly expressed during mid-gametogenesis in control cells, whereas their induction was nearly abolished following Hsf1 depletion. Consistent with these targets, Hsf1-dependent genes were enriched for functions related to protein folding, maturation, and stabilization, as well as responses to heat and oxidative stress (Fig. S3A).

**Fig. 2.**
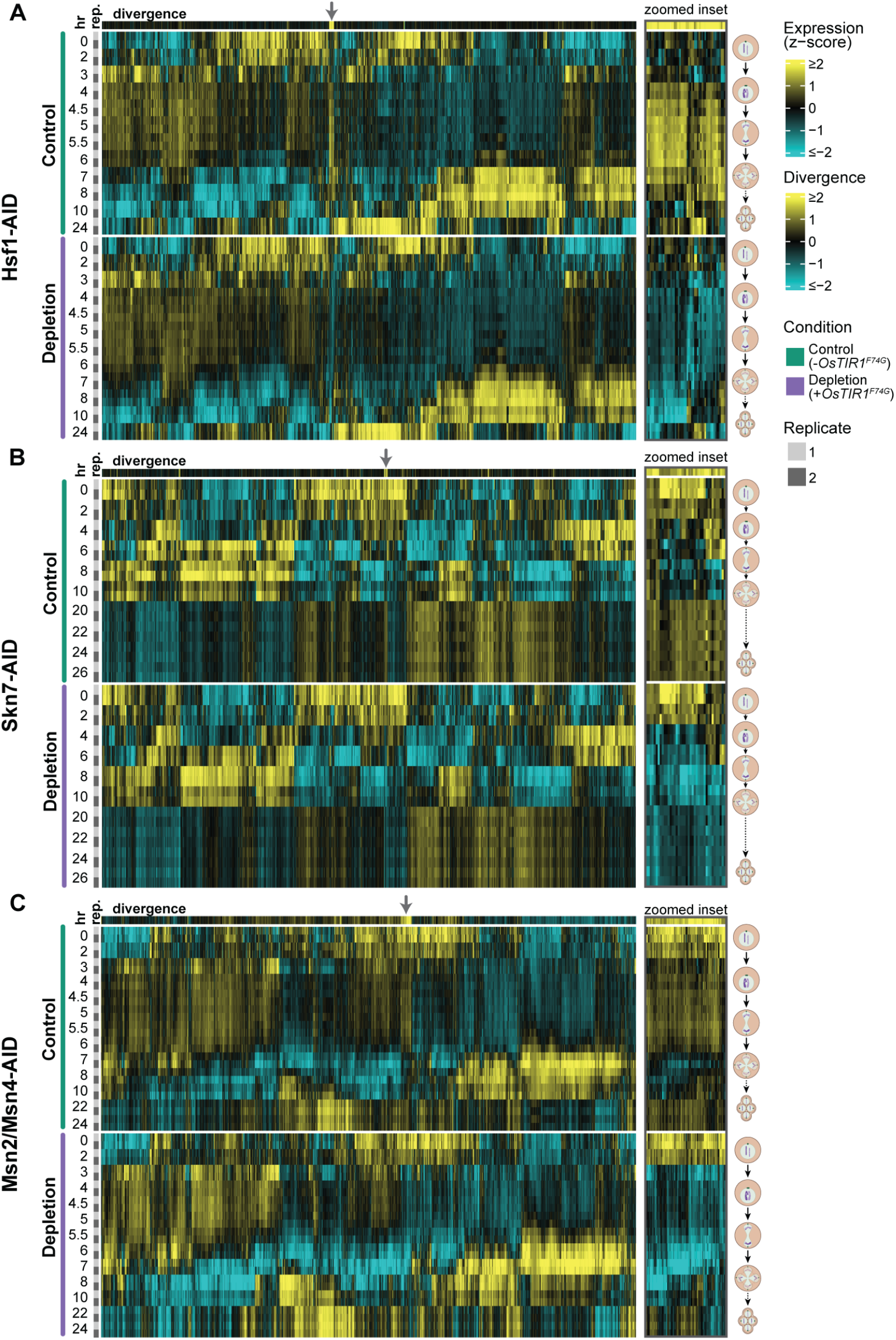
Discrete sets of genes exhibit TF-dependent expression during gametogenesis. **A-C** Heatmaps of variance-stabilized expression values (DESeq2 vst), row-scaled (z-score) across all samples. Rows represent individual samples, grouped by condition (control, depletion) and ordered by timepoint (0-26 hr in SPO) and replicate (1, 2) within each condition. Genes (columns) were hierarchically clustered using Pearson correlation distance and Ward’s linkage (ward.D2). Colors represent the number of standard deviations above (yellow) or below (cyan) each gene’s mean expression across all samples. Top row (divergence) shows the mean difference in z-scored expression (control – depletion), highlighting genes with TF-dependent expression during 5-6 hr in SPO for Hsf1 (A), 20-26 hr in SPO for Skn7 (B), and 5-6 hr in SPO for Msn2/Msn4 (C). Grey arrows indicate example clusters of genes exhibiting TF-dependent expression, and zoomed insets of these clusters are shown to the right of each heatmap.

**Fig. 3.**
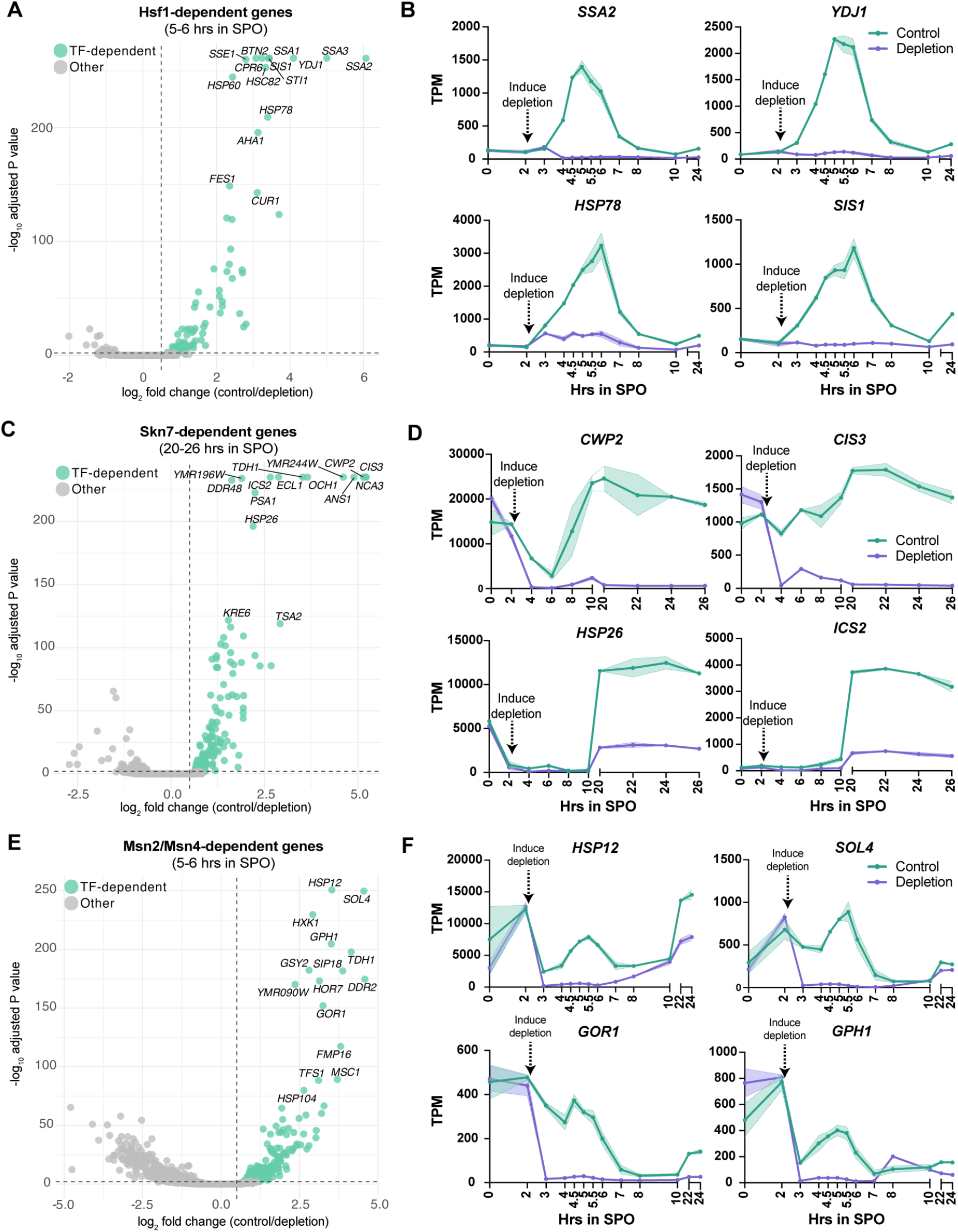
Stress-responsive TFs activate distinct transcriptional programs during gametogenesis. **A,C,E** Volcano plots showing results from differential expression analyses comparing control and depletion conditions. Samples were collected in duplicate for RNA-seq throughout a meiotic time course for Hsf1-AID (A), Skn7-AID (C), and Msn2/Msn4-AID (E) control and depletion strains, with TF depletion and synchronous meiotic entry induced at 2 hr in SPO. DESeq2 was run using only the timepoints indicated above each graph, which were the times with highest TF protein levels and activity. Dashed lines represent log_2_ fold change > 0.5 and P_adj_ < 0.01 cut-offs. Genes that met both criteria were considered TF-dependent and are shown in teal; all others are grey. The top 15 genes in each experiment are labeled. Full DESeq2 results can be found in Table S4–6. **B,D,F** Plots showing TPM levels throughout gametogenesis of 4 example TF-dependent genes identified from the Hsf1-AID (B), Skn7-AID (D), and Msn2/Msn4-AID (F) RNA-seq experiments. Shown is the average of two replicates, with the shaded region representing the range. Timepoints collected for each experiment are shown on the x-axis.

Skn7 controlled a temporally distinct transcriptional program. We profiled 10 time points spanning 0-26 hr in SPO, with increased sampling after 20 hr, coinciding with maximal Skn7 protein abundance. Control and Skn7-depleted transcriptomes remained highly correlated throughout gametogenesis (*r* ≥ 0.96) but diverged most beginning at 20 hr (Fig. 2B, S2B). Differential expression analysis across the 20-26 hr time points identified 153 Skn7-dependent genes (Fig. 3C), including *CWP2* and *CIS3*, both cell wall constituents^45–47^; *ICS2*, which is also induced by H_2_O_2_ and heat^48,49^; and *HSP26*, encoding a small heat-shock protein^50^(Fig. 3D). These genes were highly expressed during late gametogenesis in control cells but markedly reduced following Skn7 depletion. Skn7-dependent genes were enriched for functions related to cell-wall organization and biogenesis, metabolism, responses to chemical and oxidative stress, and redox homeostasis (Fig. S3B).

Msn2 and Msn4 regulated a third transcriptional program during mid-gametogenesis. We profiled 13 time points spanning 0-24 hr in SPO, with increased sampling around 5 hr, when Msn2 and Msn4 protein abundance peaks. Global transcription remained similar between control and Msn2/Msn4-depleted cells throughout most of the time course but diverged during mid-gametogenesis, with the correlation reaching a minimum of r = 0.86 at 6 hr (Fig. 2C, S2C). Differential expression analysis of the 5, 5.5, and 6 hr time points identified 274 Msn2/Msn4-dependent genes (Fig. 3E), including *HSP12*, a stress-induced plasma-membrane-associated protein^51–53^; *SOL4*, encoding 6-phosphogluconolactonase^54^; *GOR1*, encoding glyoxylate reductase^55^; and *GPH1*, encoding glycogen phosphorylase^56^ (Fig. 3F). These genes were strongly expressed during mid-gametogenesis in control cells but exhibited little to no expression following Msn2/Msn4 depletion. Msn2/Msn4-dependent genes were enriched for functions related to metabolism, cellular quality control, and oxidative-stress responses (Fig. S3C).

Together, these analyses show that Hsf1, Skn7, and Msn2/Msn4 regulate temporally structured transcriptional programs during gametogenesis. The timing of each program closely corresponds to the expression dynamics of its cognate TF, with Hsf1 and Msn2/Msn4 activity concentrated during mid-gametogenesis and Skn7 activity emerging later. Although all three programs include genes associated with cellular stress responses, they show distinct functional enrichments, suggesting that multiple stress-responsive TFs contribute to homeostatic functions at different stages of gametogenesis.

### Msn2, Msn4 and Skn7 act together in gametogenesis to promote gamete viability and fitness

The induction of multiple stress-responsive transcriptional programs during gametogenesis raised the possibility that these TFs contribute to the fitness of the resulting gametes, known as spores in yeast. To test this, we depleted Hsf1, Skn7, or Msn2/Msn4 during gametogenesis and assessed the fitness of the resulting spores. Following tetrad dissection onto nutrient-rich medium lacking CuSO_4_ and 5-Ph-IAA, allowing the depleted TFs to be resynthesized, we classified individual spores as healthy or impaired based on their ability to form normal-sized colonies (Fig. 4A). Surprisingly, depletion of Hsf1, Skn7, or Msn2/Msn4 individually did not impair spore fitness, with control and depletion conditions exhibiting similarly low frequencies of impaired spores (Fig. 4B).

**Fig. 4.**
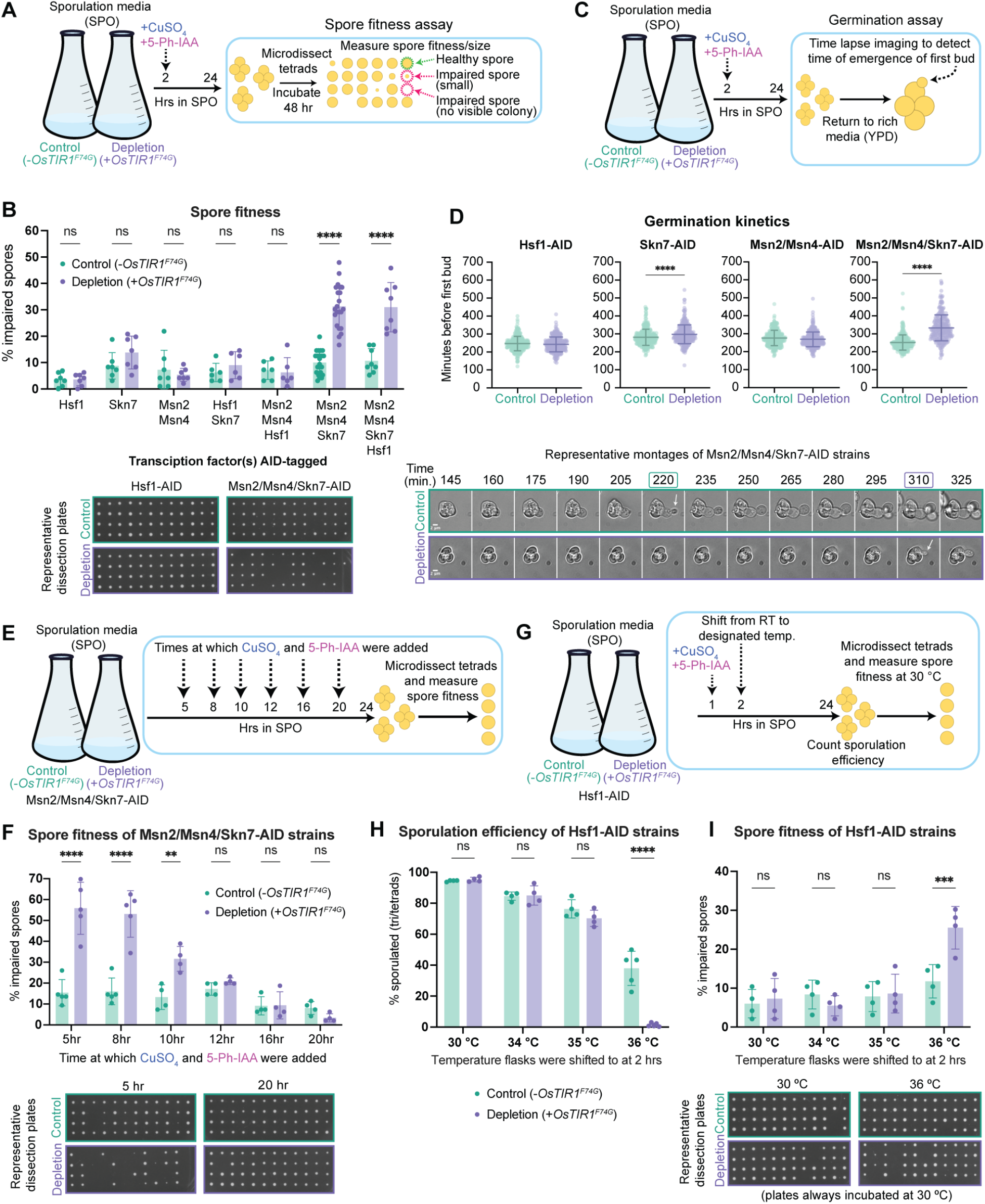
Stress-responsive TFs act during gametogenesis to promote gamete fitness. **A** Schematic showing spore fitness experiment. Diploid cells are induced to sporulate ± TF depletion at 2 hr in SPO. After 24 hr, sporulated cells are washed to remove CuSO_4_ and 5-Ph-IAA, and tetrads are separated by microdissection onto rich media plates (YPD). Plates are incubated at 30 °C for 48 hr and imaged. Spore colony size is quantified in FIJI by image thresholding and particle size analysis. Impaired spores are defined as colonies that are either not visible (indicating failure to germinate, or so slow growing that no colony is visible at 48 hr), or smaller than half the size of the average colony on the corresponding control (-*OsTIR1^F74G^*) plate. **B** Spore fitness results ± individual or combinatorial meiotic TF depletion. Shown is the average ± standard deviation (n ≥ 6). Statistical significance determined by two-way ANOVA with matched values and Tukey’s multiple comparisons correction (**** P_adj_ < 0.0001). Representative plates from Hsf1-AID depletion and Msn2/Msn4/Skn7-AID depletion experiments shown below. **C** Schematic showing germination kinetics imaging experiment. Diploid cells are induced to sporulate ± TF depletion at 2 hr in SPO. After 24 hr, sporulated cells are washed to remove CuSO_4_ and 5-Ph-IAA and loaded onto a Concanavalin A-treated 96-well plate. YPD is added and the time of emergence of the first bud is scored relative to rich media addition. Cells are imaged by wide-field microscopy every 5 minutes. **D** Germination kinetics results ± individual or combinatorial meiotic TF depletion. Shown is the average ± standard deviation, with each point representing one cell (n = 3 experiments with ≥ 100 cells per condition in each experiment). Statistical significance determined by unpaired two-tailed t-test (**** P < 0.0001). Representative montages from Msn2/Msn4/Skn7-AID depletion experiment shown below. Timepoint of first bud emergence in control condition is boxed in teal, and in depletion condition is boxed in purple. White arrows point to first bud, scale bar indicates 2 µm. **E** Schematic of spore fitness experiment following varying Msn2/Msn4/Skn7-AID depletion time. Diploid cells are induced to sporulate, and depletion is induced at various timepoints. As in (A) spore fitness is measured by washing cells, dissecting tetrads on YPD, and quantifying spore colony size. **F** Spore fitness results ± Msn2/Msn4/Skn7-AID depletion at different timepoints. Shown is the average ± standard deviation (n ≥ 4). Statistical significance determined by two-way ANOVA with matched values and Tukey’s multiple comparisons correction (**** P_adj_ < 0.0001, ** P_adj_ < 0.005). Representative plates from 5-and 20-hr depletion experiments shown below. **G** Schematic of heat shock gametogenesis experiment. Diploid cells are induced to sporulate at room temperature, Hsf1-AID depletion is induced at 1 hr, and cultures are shifted to the designated temperature at 2 hr. After 24 hr, sporulation efficiency is measured, and as in (A), spore fitness is measured by washing cells, dissecting tetrads on YPD, incubating plates at 30 °C, and quantifying spore colony size. **H** Sporulation efficiency results from Hsf1-AID meiotic depletion at various temperatures. Shown is the average ± standard deviation (n ≥ 4). Statistical significance determined by two-way ANOVA with matched values and Tukey’s multiple comparisons correction (**** P_adj_ < 0.0001). **I** Spore fitness results from Hsf1-AID meiotic depletion at various temperatures. Shown is the average ± standard deviation (n = 4). Statistical significance determined by two-way ANOVA with matched values and Tukey’s multiple comparisons correction (*** P_adj_ < 0.0005). Representative plates from 30 °C and 36 °C experiments shown below.

Because these TFs regulate some overlapping homeostatic functions, we asked whether functional redundancy might mask phenotypes following individual depletion. Co-depletion of Hsf1/Skn7 or Hsf1/Msn2/Msn4 did not significantly impair spore fitness. In contrast, simultaneous depletion of Msn2, Msn4, and Skn7 increased the fraction of impaired spores from 10% in control cells to 31% following depletion (Fig. 4B). Addition of Hsf1 depletion did not further exacerbate this phenotype, with 31% impaired spores following depletion of all four TFs (Fig. 4B). Robust and sustained depletion of all four proteins was maintained in the combined AID strain (Fig. S4A), and all strains sporulated with high efficiency (Fig. S4B). We therefore focused subsequent experiments on Msn2/Msn4/Skn7 depletion, the minimal combination sufficient to impair spore fitness.

To assess another feature of gamete health, we measured the kinetics of spore germination following meiotic TF depletion. Spores remain dormant until exposed to nutrient-rich conditions, after which they germinate and resume mitotic growth. Using time-lapse imaging, we measured the interval between addition of rich medium and emergence of the first bud^57^ (Fig. 4C). Meiotic depletion of Hsf1 or Msn2/Msn4 did not alter germination timing, whereas Skn7 depletion produced a modest but significant 17-min delay (Fig. 4D). Simultaneous depletion of Msn2/Msn4/Skn7 caused a substantially larger 81-min delay relative to the corresponding control. Together, the spore-fitness and germination phenotypes indicate that Msn2, Msn4, and Skn7 work together during gametogenesis to promote the production of fit gametes capable of efficiently resuming growth.

We next sought to define when Msn2/Msn4/Skn7 activity is required to promote gamete fitness. Whereas TF depletion in previous experiments was initiated 2 hr after transfer to SPO, we induced depletion at progressively later time points during gametogenesis (Fig. 4E). Depletion initiated at 5, 8, or 10 hr resulted in 56%, 53%, and 32% impaired spores, respectively, whereas depletion initiated at 12, 16, or 20 hr did not significantly impair spore fitness relative to the corresponding controls (Fig. 4F). TF depletion remained robust following late induction, indicating that the absence of a phenotype was not due to inefficient degradation (Fig. S4C), and both strains sporulated well across all depletion timepoints tested (Fig. S4D). Thus, Msn2/Msn4/Skn7 activity is required during an early-to-mid-gametogenesis window to establish gamete fitness, whereas depletion initiated at 12 hr or later no longer compromises this outcome.

Because gametogenesis resets replicative lifespan in budding yeast regardless of progenitor-cell age^58^, we also asked whether Msn2/Msn4/Skn7 contribute to this aspect of gamete rejuvenation. Using an established microfluidic assay^59^, we measured the replicative lifespan of spores generated from young or aged progenitor cells following control or Msn2/Msn4/Skn7-depletion conditions (Fig. S5A–C). Among spores successfully recovered for analysis, meiotic TF depletion did not significantly alter replicative lifespan in progeny derived from either young or aged cells (Fig. S5C). Because impaired spores that fail to germinate efficiently may be underrepresented during microfluidic loading, we cannot exclude an effect within this population. Nevertheless, among successfully germinating spores, Msn2/Msn4/Skn7 depletion does not detectably disrupt meiotic lifespan resetting.

### Hsf1 promotes successful gametogenesis at elevated temperature

The absence of an overt gamete-fitness phenotype following Hsf1 depletion was unexpected given its essential cellular functions and the reported roles of heat-shock factors during gametogenesis in other organisms^60–62^. Because gametogenesis is temperature-sensitive across species, including budding yeast^63–66^, we asked whether Hsf1 becomes functionally important during gametogenesis at elevated temperature (Fig. 4G).

At 30 °C, Hsf1 control and depletion strains sporulated efficiently, producing 94% and 95% triads or tetrads, respectively (Fig. 4H). Sporulation efficiency decreased at 34 °C and 35 °C but remained similar between control and Hsf1-depleted cells. At 36 °C, however, sporulation efficiency declined sharply and was substantially lower following Hsf1 depletion, with only 1% of Hsf1-depleted cells forming triads or tetrads compared with 38% of control cells (Fig. 4H).

Hsf1 depletion also compromised the fitness of the spores that were successfully produced at 36 °C. Whereas depletion at 30 °C, 34 °C, or 35 °C did not significantly increase the fraction of impaired spores, depletion at 36 °C resulted in 26% impaired spores compared with 12% in the corresponding control (Fig. 4I). Thus, although Hsf1 is not limiting for gamete production or fitness under standard meiotic conditions, it becomes important for both efficient sporulation and production of fit gametes when gametogenesis occurs at elevated temperature.

### Msn2, Msn4, and Skn7 promote gamete fitness through antioxidant gene expression

The finding that simultaneous depletion of Msn2/Msn4/Skn7 impaired gamete fitness, whereas depletion of Msn2/Msn4 or Skn7 alone did not, suggested that these TFs redundantly regulate one or more genes required for gamete fitness. To identify these genes, we performed RNA-seq across a meiotic time course following simultaneous Msn2/Msn4/Skn7 depletion (Fig. 5A). We focused our analysis on 5-6 hr in SPO, when Msn2/Msn4 abundance is high and when depletion initiated during this period produced the strongest spore-fitness phenotype (Fig. 4F). Differential expression analysis comparing control and depletion conditions identified 201 Msn2/Msn4/Skn7-dependent genes (Fig. 5B), including *HSP12*, *SOL4*, *DDR2*, and *FMP16*, which were also identified as Msn2/Msn4-dependent (Fig. 3E), as well as *NCA3*, which was also identified as Skn7-dependent (Fig. 3C).

**Fig. 5.**
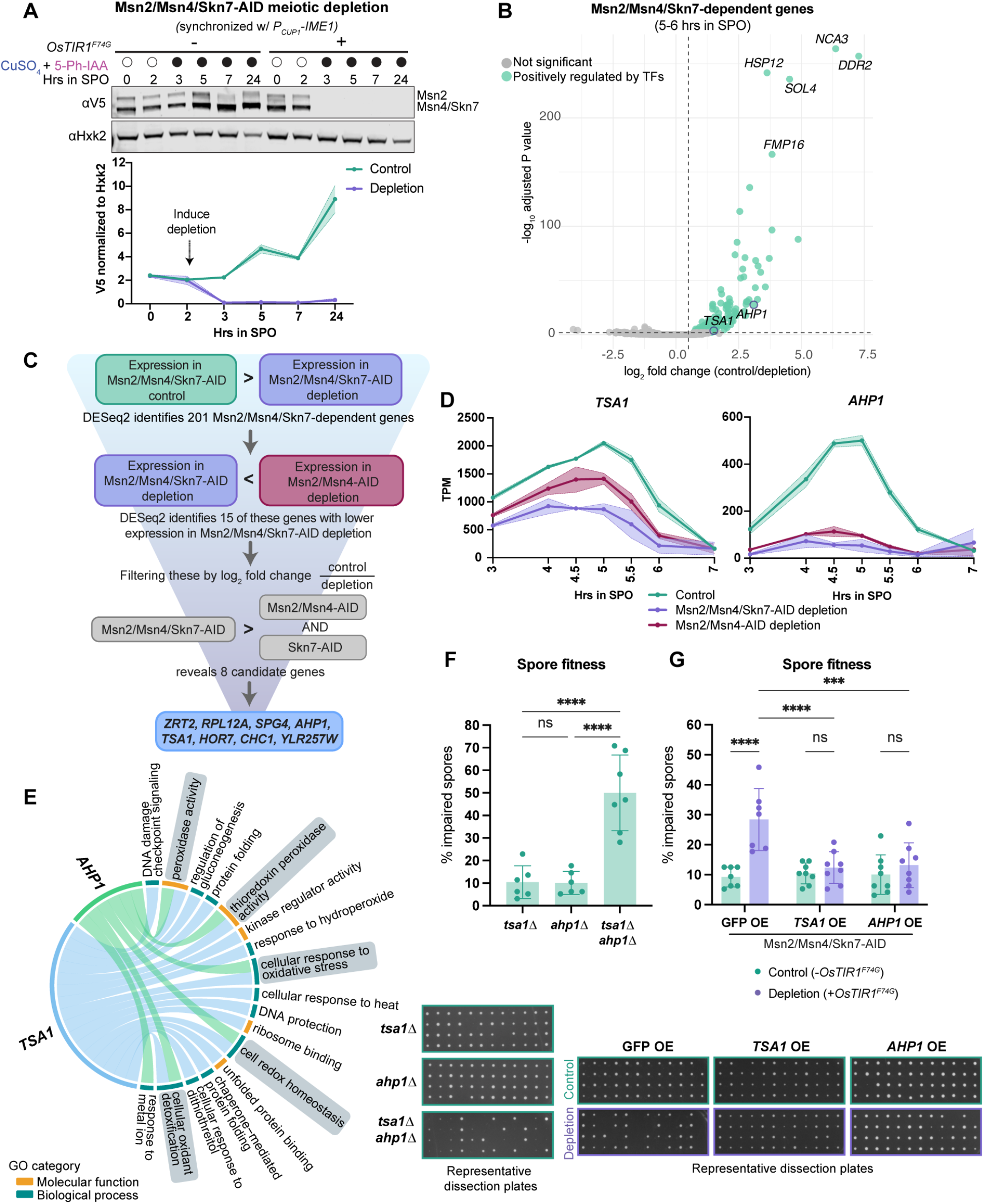
Msn2, Msn4, and Skn7 promote gamete fitness through antioxidant gene expression. **A** Western blot showing Msn2/Msn4/Skn7-AID levels throughout gametogenesis in control (-*OsTIR1^F74G^*) and depletion (+*OsTIR1^F74G^*) strains. Msn4-AID and Skn7-AID are only 1 kDa apart in size and are indistinguishable on the blot. Depletion and synchronous meiotic entry were induced at 2 hr in SPO by addition of 50 µM CuSO_4_ and 100 µM 5-Ph-IAA. Hxk2 serves as a loading control. Quantification below blot, shown is the average of two replicates, with shaded region representing the range. **B** Volcano plot showing results from differential expression analysis comparing Msn2/Msn4/Skn7-AID control and depletion conditions. Samples were collected in duplicate for RNA-seq throughout a meiotic time course with depletion and synchronous meiotic entry induced at 2 hr in SPO. DESeq2 was run using only the timepoints indicated above the graph. Dashed lines represent log_2_ fold change > 0.5 and P_adj_ < 0.01 cut-offs. Genes that met both criteria were considered TF-dependent and are shown in teal; all others are grey. The top 5 genes are labeled, in addition to *AHP1* and *TSA1*, which are circled in blue. Full DESeq2 results can be found in Table S7. **C** Flow chart demonstrating how genes disproportionally affected by Msn2/Msn4/Skn7-AID depletion relative to Msn2/Msn4-AID and Skn7-AID depletion were identified. See methods for details. **D** Plots showing TPM levels during gametogenesis of *TSA1* and *AHP1*. Shown is the average of two replicates, with shaded region representing the range. Control is from the Msn2/Msn4/Skn7-AID experiment. **E** Chord diagram linking *TSA1* (blue) and *AHP1* (green) to their annotated GO molecular function (orange) and biological process (dark teal) terms. Ribbon width is uniform (binary presence/absence of each annotation per gene). Shared annotations are shaded in grey. **F** Results from spore fitness experiments with deletion strains in Msn2/Msn4-AID control background. Experiment conducted as in Fig. 4A. Shown is the average ± standard deviation (n ≥ 6). Statistical significance determined by one-way ANOVA with Tukey’s multiple comparisons correction (**** P_adj_ < 0.0001). Representative dissection plates from each genotype shown below. **G** Results from spore fitness experiments with meiotic overexpression strains in Msn2/Msn4/Skn7-AID control and depletion backgrounds. Experiment conducted as in Fig. 4A. Shown is the average ± standard deviation (n ≥ 7). Statistical significance determined by two-way ANOVA with matched values and Tukey’s multiple comparisons correction (**** P_adj_ < 0.0001, *** P_adj_ < 0.0005). Representative dissection plates from each genotype shown below.

We reasoned that genes disproportionately affected by simultaneous Msn2/Msn4/Skn7 depletion relative to depletion of Msn2/Msn4 or Skn7 alone might account for the spore-fitness phenotype. Globally, however, the Msn2/Msn4/Skn7-and Msn2/Msn4-depleted transcriptomes were remarkably similar, with Pearson correlation coefficients ≥ 0.98 at all timepoints (Fig. S6A). We therefore searched for more subtle differences among TF-dependent genes. Of the 201 Msn2/Msn4/Skn7-dependent genes, 15 exhibited significantly lower expression following Msn2/Msn4/Skn7 depletion than following Msn2/Msn4 depletion during mid-gametogenesis. We further prioritized genes whose dependence on the combined TF depletion exceeded that observed following either Msn2/Msn4 or Skn7 depletion alone. This analysis yielded eight candidate genes: *ZRT2, RPL12A, SPG4, AHP1, TSA1, HOR7, CHC1,* and *YLR257W* (Fig. 5C,D; Fig. S6B).

Among these candidates, we noted that two, *TSA1* and *AHP1*, are both antioxidant genes with shared functions. *TSA1* encodes a thioredoxin peroxidase that protects the cytosol and translating ribosomes from reactive oxygen species and can also function as a molecular chaperone under oxidative stress^67–69^; *AHP1* also encodes a thioredoxin peroxidase with overlapping antioxidant functions with Tsa1^70,71^ (Fig. 5E). We asked whether loss of either or both genes could recapitulate the spore-fitness defect caused by Msn2/Msn4/Skn7 depletion. Deletion of *TSA1* or *AHP1* alone did not substantially impair spore fitness, both resulting in 10% impaired spores (Fig. 5F). Strikingly, however, simultaneous deletion of *TSA1* and *AHP1* resulted in 52% impaired spores (Fig. 5F). This closely phenocopies the most severe 56% defect caused by Msn2/Msn4/Skn7 depletion at 5 hr (Fig. 4F) and demonstrates that loss of these two genes alone is sufficient to compromise gamete fitness. Deleting one or both genes in the Msn2/Msn4-AID depletion background did not exacerbate this phenotype (Fig. S7A), suggesting additional Msn2/Msn4-dependent genes do not play a significant role in promoting spore fitness. Both TFs remained robustly depleted in these strains (Fig. S7B–D).

We next asked whether restoring expression of either thioredoxin peroxidase could bypass the requirement for Msn2/Msn4/Skn7. We replaced the endogenous promoter of either *TSA1* or *AHP1* with the *P_ATG8_* promoter, which drives high expression throughout gametogenesis^6^, and assessed spore fitness following Msn2/Msn4/Skn7 depletion. Remarkably, meiotic expression of either *P_ATG8_*-*TSA1* or *P_ATG8_*-*AHP1* was sufficient to rescue the spore-fitness defect despite near-complete depletion of all three TFs (Fig. 5G; Fig. S7E,F). In contrast, expression of *P_ATG8_*-*GFP* did not rescue the phenotype, resulting in 28% impaired spores following TF depletion (Fig. 5G; Fig. S7G) and closely resembling the 31% impaired spores seen following Msn2/Msn4/Skn7 depletion without any overexpression allele (Fig. 4B).

Together, these loss-and gain-of-function experiments identify the thioredoxin peroxidases Tsa1 and Ahp1 as critical downstream effectors of Msn2, Msn4, and Skn7 during gametogenesis. Loss of both antioxidant enzymes is sufficient to phenocopy TF depletion, whereas restoring expression of either enzyme is sufficient to bypass the requirement for these TFs in promoting spore fitness. These results identify antioxidant gene expression as a critical physiological output of Msn2, Msn4, and Skn7 during gametogenesis and reveal a potential means by which endogenous activation of stress-responsive transcriptional programs promote the fitness of the resulting gametes.

## Discussion

Stress-response TFs are traditionally studied for their roles in protecting cells from environmental perturbations. However, the cellular processes they regulate, including proteostasis and redox homeostasis, are also critical under physiological conditions. Here, we identify gametogenesis as an endogenous developmental context in which four stress-response TFs–Hsf1, Skn7, Msn2, and Msn4–regulate gene expression in the absence of imposed environmental stress. We show that combined depletion of Msn2, Msn4, and Skn7 during gametogenesis compromises the fitness of the resulting spores and delays their germination. Remarkably, we identify two peroxiredoxin genes, *TSA1* and *AHP1*, whose loss impairs spore fitness and whose overexpression rescues the defects caused by TF depletion. These findings establish a physiological role for stress-response transcriptional programs during gametogenesis and demonstrate that the expression of a small subset of antioxidant genes can have a substantial impact on the fitness of meiotic progeny.

### Stress-response transcription factors contribute to the meiotic transcriptional program

Meiotic differentiation requires extensive transcriptional reprogramming to coordinate chromosome segregation, cellular remodeling, and gamete formation^4–6,72^. In budding yeast, the core transcriptional circuitry controlling meiotic progression has been extensively characterized^1,5,27,73–76^. Our findings expand this regulatory framework by identifying genes whose meiotic expression depends on TFs traditionally associated with environmental stress responses. Although these genes include many established stress-response targets, their regulation during gametogenesis suggests that canonical stress-response transcriptional programs are also integrated into physiological developmental processes.

Indeed, we observed substantial overlap between genes dependent on these TFs during gametogenesis and those previously identified under vegetative growth or stress conditions. All 18 Hsf1-dependent genes reported by Solís et al.^32^ were also Hsf1-dependent during gametogenesis. Similarly, approximately half of the previously identified Msn2/Msn4-dependent genes^29,33,40^ and 10 of the 14 previously reported Skn7-dependent genes^18,39,77,78^ were dependent on their respective TFs during gametogenesis. Consistent with these overlaps, the meiotic TF-dependent gene sets were enriched for stress-response functions. Thus, rather than activating entirely distinct transcriptional programs, meiotic cells appear to deploy components of established stress-response networks during differentiation.

How these TFs are integrated into the meiotic transcriptional program remains an important question. Their activity could be driven by developmental signals that engage canonical stress-response pathways or by changes in cellular physiology intrinsic to meiotic differentiation. Furthermore, the extent to which the observed transcriptional changes reflect direct TF regulation remains to be determined. Nevertheless, our findings suggest that meiotic gene regulation extends beyond dedicated developmental regulators to include homeostatic transcriptional networks that support the physiological demands of differentiation.

### Distinct contributions of stress-response TFs to gamete fitness

Although all four TFs tested here regulate gene expression during gametogenesis, their contributions to gamete fitness differ. This distinction was particularly striking for Hsf1. Heat shock factors have established roles in germ cell development and gametogenesis across several animal species, including *Drosophila melanogaster*, *Caenorhabditis elegans*, *Danio rerio*, and *Mus musculus*^61,62^. In budding yeast, however, the role of Hsf1 during gametogenesis has not previously been characterized. Given its essential function during vegetative growth and the importance of HSFs for reproduction in other organisms, we anticipated that meiotic Hsf1 depletion would compromise spore formation or fitness. Surprisingly, we observed no obvious spore fitness defect under standard sporulation conditions, despite the dependence of numerous meiotic genes on Hsf1.

At elevated temperature (36 °C), however, Hsf1 depletion adversely affected sporulation efficiency and spore fitness. One possibility is that Hsf1-dependent gene expression during gametogenesis provides a protective capacity that becomes particularly important under challenging conditions. By promoting the expression of chaperones and other proteostasis factors, Hsf1 may help meiotic cells accommodate fluctuations in temperature or increased demands on protein homeostasis. This function could be particularly relevant given the temperature sensitivity of gametogenesis relative to mitotic growth^63–66^. Alternatively, Hsf1 may contribute to aspects of spore physiology that are not captured by our assays under standard conditions. Thus, while Hsf1 regulates a transcriptional program during gametogenesis, its contribution to spore fitness appears to be condition dependent.

In contrast, combined depletion of Msn2, Msn4, and Skn7 impaired spore fitness under standard conditions, revealing a requirement for these regulators even in the absence of imposed environmental stress. This phenotype was particularly striking because the transcriptional profiles of Msn2/Msn4-depleted and Msn2/Msn4/Skn7-depleted cells were remarkably similar. Only eight genes exhibited significantly lower expression upon additional depletion of Skn7, yet this relatively modest transcriptional difference was associated with a pronounced spore fitness defect. These observations show that the functional consequences of disrupting a transcriptional network are not necessarily proportional to the extent of global gene expression changes. Rather, a small number of critical transcriptional outputs may have a disproportionate impact on developmental fitness without obvious secondary effects on gene expression. Indeed, we found that two of these genes, *TSA1* and *AHP1*, are sufficient to account for key aspects of the observed phenotype, highlighting the importance of specific downstream effectors within a broader transcriptional program.

### Peroxiredoxin expression supports the fitness of meiotic progeny

Among the genes regulated by Msn2, Msn4, and Skn7, we identified *TSA1* and *AHP1* as particularly important for spore fitness. Loss of these genes impaired spore fitness, whereas overexpression of either gene rescued the defects caused by combined TF depletion. These findings establish a functional connection between stress-response TF activity, peroxiredoxin expression, and the production of healthy meiotic progeny. Although the broader Msn2/Msn4/Skn7-dependent transcriptional program comprises more than 200 genes, restoring the expression of individual antioxidant components was sufficient to compensate for the loss of multiple upstream regulators in our spore fitness assays.

Tsa1 and Ahp1 are members of the peroxiredoxin family, which contributes to peroxide detoxification and redox homeostasis through the thioredoxin system^70,79^. Notably, we also identified *TSA2*, encoding the third cytosolic peroxiredoxin in budding yeast, as an Msn2/Msn4/Skn7-dependent gene during gametogenesis (Fig. 5B; Table S7). The coordinated regulation of all three cytosolic peroxiredoxins suggests that maintenance of redox homeostasis may be an important function of this transcriptional network during gamete formation. Although Tsa1 is considered the dominant peroxide-scavenging enzyme in budding yeast^70,80^, overexpression of either *TSA1* or *AHP1* rescued the spore fitness defect caused by TF depletion. This functional compensation is consistent with previous observations that increased Ahp1 expression can restore peroxide-scavenging capacity in cells lacking Tsa1^70^.

Why might peroxiredoxin expression be particularly important during gametogenesis? Although oxidative stress is commonly studied through exposure to exogenous oxidants, reactive oxygen species are also continuously generated as byproducts of normal cellular processes, including aerobic metabolism, which is required for gametogenesis^14,19^. Meiotic differentiation involves extensive physiological remodeling, including dramatic rearrangements of the mitochondrial and ER networks^9,10^, and developing gametes must maintain cellular integrity while undergoing major changes in cellular organization and metabolism. We therefore propose that Msn2/Msn4/Skn7-dependent peroxiredoxin expression helps maintain redox homeostasis during this transition, supporting the fitness of the resulting spores. Although this model is consistent with the established antioxidant functions of Tsa1 and Ahp1, whether their meiotic requirement reflects peroxide detoxification, redox signaling, or other functions of peroxiredoxins remains to be determined.

An intriguing possibility is that the activation of antioxidant genes during gametogenesis serves not only to protect cells as they differentiate but also to prepare the resulting gametes for subsequent challenges. Because TF depletion during gametogenesis compromises spore fitness and delays germination, the transcriptional program established during gamete formation may influence the physiological competence of the resulting progeny. Peroxiredoxins expressed during gametogenesis could protect differentiating cells from endogenous oxidative damage, establish an antioxidant capacity that is inherited by newly formed spores, or support the transition from dormancy to growth during germination. These possibilities are not mutually exclusive and raise the broader question of how transcriptional programs activated during differentiation influence cellular fitness beyond the developmental transition itself.

The importance of redox homeostasis during gamete formation extends beyond budding yeast. Oxidative stress has been implicated in impaired germ cell development and reproductive function across diverse organisms, including humans^81^. Moreover, the thioredoxin system has established roles in germ cell biology, and specialized thioredoxins have been identified in the reproductive tissues of both invertebrates and mammals^82–87^. Our findings add to this broader picture by demonstrating that transcriptional regulation of peroxiredoxins during gametogenesis is important for the fitness of yeast gametes. Although the specific regulatory mechanisms may differ across species, maintaining redox homeostasis during gamete development may represent a shared physiological requirement.

More broadly, our study highlights how transcriptional programs traditionally associated with environmental stress can serve essential functions during normal development. Rather than functioning exclusively as reactive defenses against external perturbations, stress-response networks can be integrated into developmental programs to support cellular homeostasis and differentiation. In budding yeast gametogenesis, this includes the coordinated regulation of antioxidant genes that promote the fitness of meiotic progeny. Together, our findings reveal that stress-response programs are not merely activated in response to environmental challenges but can be deployed as part of developmental programs to support the fitness of the next generation.

## Author contributions

A.L.B., E.Ü., and G.A.B. designed the research. A.L.B., S.S., and D.Y. performed the experiments. A.L.B., S.S., and D.Y. analyzed the data. A.L.B. and E.Ü. wrote the original draft of the manuscript. All authors revised the final manuscript.

## Acknowledgements

We thank Nicholas Ingolia, Michael Rapé, Craig Miller, Jeremy Thorner, Benjamin Styler, Thomas Moss, Tianyao Xiao, Alejandro Collins, Him Visitvoranat, Jessica Leslie, Silvan Spiri, Claudia Medrano, Cyrus Ruediger, and the entire Brar-Ünal lab for comments on this manuscript and for valuable discussions.

## Materials and Methods

### Strain generation

All experiments were performed using *Saccharomyces cerevisiae* strains in the SK1 background. All yeast strains, plasmids, and primers used in this study are listed in Table S1, Table S2, and Table S3, respectively. Constructs were generated using NEBuilder HiFi DNA Assembly Master Mix (New England Biolabs, E2621) and sequenced verified using whole plasmid sequencing. Deletions, endogenous C-terminal tagging, promoter swaps, and MX cassette swaps were performed using standard PCR-based methods^82,83^. Yeast transformation was performed using the lithium acetate method^84^, and proper integration was confirmed by PCR, sequencing, and Western blot (where applicable). Transformants were backcrossed prior to subsequent use.

The following alleles were generated in previous studies: P*_CUP1_-*OsTIR1^F74G^^85^; P*_CUP1_-IME1*^28^; *ime1Δ*^86^; P*_PMA2_-PMA2*-GFP, P_tetO_*-STE5*, P*_RNR2_*-TetR-*TUP1*, and P_tetO_-TetR, *flo8Δ*, *amn1^BY4741^* ^59^.

The following alleles were generated in this study: *HSF1*-3V5-IAA17 (*HSF1*-AID), *SKN7*-3V5-IAA17 (*SKN7*-AID), *MSN2*-3V5-IAA17 (*MSN2*-AID), *MSN4*-3V5-IAA17 (*MSN4*-AID), *tsa1Δ*, *ahp1Δ*, P*_ATG8_-TSA1*, P*_ATG8_-AHP1*, P*_ATG8_*-GFP, and various MX cassette marker swaps of these alleles.

The C-terminal 3V5-IAA17 (AID) tags were generated by inserting the tag and a KanMX cassette at the endogenous locus for each transcription factor, replacing the stop codon. The *tsa1Δ* and *ahp1Δ* deletions were generated by replacing the entire coding sequence of each gene with a NatMX cassette. The P*_ATG8_-TSA1* and P*_ATG8_-AHP1* alleles were generated by inserting a HygMX cassette and P*_ATG8_* promoter directly upstream of each endogenous coding sequence. The P*_ATG8_-*GFP allele was generated by inserting a HygMX and P*_ATG8_-GFP* sequence (pÜB3117 digested with NotI) at the *HO* locus (chosen as a neutral genomic location).

### Sporulation conditions

Yeast strains were thawed on YPG (1% yeast extract, 2% peptone, 3% glycerol, 2% agar) plates and grown at 30 °C overnight. The next morning, strains were transferred to YPD 4% (1%yeast extract, 2% peptone, 4% glucose, 2% agar) for further outgrowth. Cells were then grown in liquid YPD (1% yeast extract, 2% peptone, 2% glucose, 22.4 mg/L uracil, and 80 mg/L tryptophan) at 30 °C on a shaker for approximately 20 hr, reaching saturation (OD_600_ ≥ 10). Cultures were then back diluted to OD_600_ = 0.25 in BYTA (1% yeast extract, 2% bacto tryptone, 1% potassium acetate, and 50 mM potassium phthalate) and grown at 30 °C on a shaker for another 20 hr, reaching OD_600_ ≥ 5. To induce sporulation, cells were pelleted by centrifugation for 2 minutes at 2,000 rcf at RT, washed with MilliQ H_2_O, and resuspended in SPO (2% potassium acetate with 0.04 g/L adenine, 0.04 g/L uracil, 0.01 g/L histidine, 0.01 g/L leucine and 0.01 g/L tryptophan, adjusted to a pH of 7 with acetic acid) at OD_600_ = 1.9. Cultures were maintained at 30 °C on a shaker for 24 hr. Throughout the procedure, flasks with a volume ≥ 10x the culture volume were used to maintain proper aeration. For experiments using the P*_CUP1_-IME1* synchronization system^28^, cells were arrested prior to meiotic entry due to lack of *IME1* expression, and released into gametogenesis upon addition of 50 µM CuSO_4_ at 2 hr in SPO.

For all depletion experiments, we used the AID2 system^26^, which exhibits reduced basal degradation compared to the original AID system^87^. The F74G mutated *OsTIR1* (*Os*; *Oryza sativa*) gene was placed under the control of the copper-inducible P*_CUP1_* promoter. To induce depletion, 50 µM CuSO_4_ (Sigma Aldrich, 209198) and 100 µM of the auxin analog 5-Ph-IAA (MedChemExpress, HY-134653) were added to SPO cultures. All control and depletion strains were histidine prototrophy matched.

### Immunoblotting

To prepare samples for immunoblotting, trichloroacetic acid (TCA) extraction was performed. 4 OD_600_ units of cells were centrifuged for 2 minutes at 2,000 rcf at 4 °C. Media was aspirated and cells were resuspended in 1 mL of 5% TCA (Fisher Scientific, SA433-500) and incubated at 4 °C for ≥ 20 minutes before being centrifuged for 2 minutes at 1,000 rcf at RT. Supernatant was aspirated, and cell pellets were washed with 1 mL of acetone. Samples were centrifuged for 5 minutes at 20,000 rcf at RT and acetone was carefully aspirated. Pellets were dried in a fume hood overnight. The next day, pellets were resuspended in 100 µL protein breakage buffer (50 mM Tris pH 7.5, 1 mM EDTA, 3 mM DTT, 1X Roche cOmplete EDTA-free inhibitor cocktail) and approximately 100 µL acid-treated glass beads (Sigma Aldrich, G8772), then placed in a Mini-Beadbeater-96 (BioSpec) for 5 minutes to lyse cells. 50 µL of 3X SDS sample buffer (250 mM Tris pH 6.8, 8% β-mercaptoethanol, 40% glycerol, 12% SDS, 0.00067% bromophenol blue) was then added to each sample. Samples were then denatured at 95 °C for 5 minutes and centrifuged for 5 minutes at 20,000 rcf at RT before electrophoresis.

For immunoblotting, 6 µL of each sample was loaded onto a 4–12% Bis-Tris Bolt gel (Thermo Fisher) and run at 150V for 55 minutes in 1X Bolt MES SDS running buffer (Thermo Fisher, B000202) alongside Precision Plus Protein Dual Color Standards (BioRad, 1610394) to determine protein size. Proteins were then transferred to a 0.45 µm nitrocellulose membrane using a Trans-Blot Turbo System (Bio-Rad) with the 30-minute standard protocol. Membranes were blocked in PBS Intercept Blocking Buffer (LI-COR Biosciences, 927-70001) for 1 hr, then incubated in primary antibody diluted in PBS Intercept Blocking Buffer overnight. The following primary antibodies were used: 1:2,000 mouse anti-V5 (Invitrogen, R960-25, RRID:AB_2556564); 1:20,000 rabbit anti-hexokinase (Rockland, 100-4159, RRID:AB_219918). Membranes were washed 3 times in PBS-T (PBS with 0.1% Tween-20) and incubated in secondary antibody diluted in PBS Intercept Blocking Buffer for 2 hr at RT. The following secondary antibodies were used: 1:15,000 goat anti-mouse conjugated to IRDye 800CW (LI-COR Biosciences, 926–32212, RRID:AB_621847) and 1:15,000 goat anti-rabbit conjugated to IRDye 680RD (LI-COR Biosciences, 926–68071, RRID:AB_10956166). Blots were then washed 3 times in PBS-T and imaged on an Odyssey CLx system (LI-COR Biosciences). Image analysis and quantification was performed using ImageStudio (LI-COR Biosciences, version 5.5.4).

### RNA extraction

For all RNA extractions, cells were first collected by centrifugation for 2 minutes at 2,000 rcf at 4 °C. Media was then aspirated, and cell pellets were flash frozen in liquid N_2_. Cell pellets were thawed on ice and resuspended in 400 µL TES buffer (10mM Tris pH 7.5, 10mM EDTA, 0.5% SDS). 400 µL acid-phenol (Sigma Aldrich, P4682) and approximately 200 µL of acid-treated glass beads (Sigma Aldrich, G8772) were then added to each cell suspension. Samples were placed in a thermomixer at 65 °C and 1400 rpm for 30 minutes. Samples were then centrifuged for 5 minutes at 20,000 rcf at 4 °C, and the top aqueous layer was transferred to a new microcentrifuge tube containing 350 µL chloroform. Samples were then centrifuged again for 5 minutes at 20,000 rcf at RT, and the top aqueous layer was transferred to a low-adhesion microcentrifuge tube containing 400 µL isopropanol and 55 µL NaOAc pH 5.5. Tubes were inverted several times and incubated overnight at -20 °C. The next day, RNA was pelleted by centrifuging samples for 20 minutes at 20,000 rcf at 4 °C. RNA pellets were washed with 1 mL 80% EtOH and centrifuged again for 10 minutes at 20,000 rcf at 4 °C. EtOH was aspirated and dried RNA pellets were resuspended in DEPC-treated H_2_O. Total RNA concentrations were quantified by Qubit 3 Fluorometer (Invitrogen).

### mRNA-seq library preparation and sequencing

For all mRNA-seq experiments, samples were collected in duplicate from control and depletion strains throughout a meiotic time course. For Hsf1-AID, the following timepoints were collected: 0, 2, 3, 4, 4.5, 5, 5.5, 6, 7, 8, 10, 24 hr in SPO; for Skn7-AID, the following timepoints were collected: 0, 2, 4, 6, 8, 10, 20, 22, 24, 26 hr in SPO; for Msn2/Msn4-AID, the following timepoints were collected: 0, 2, 3, 4, 4.5, 5, 5.5, 6, 7, 8, 10, 22, 24 hr in SPO; for Msn2/Msn4/Skn7-AID, the following timepoints were collected: 0, 2, 3, 4, 4.5, 5, 5.5, 6, 7, 8, 10, 20, 22, 24, 26 hr in SPO.

Following total RNA extraction, mRNA was selected using NEXTFLEX Poly(A) Beads 2.0 (Revvity, NOVA-512993) following kit instructions. 5 µg of total RNA was used as input for all libraries. mRNA-seq libraries were then generated using the NEXTFLEX Rapid Directional RNA-Seq Kit 2.0 (Revvity, NOVA-5198-03), following kit instructions. NEXTFLEX Unique Dual Index Barcodes (Revvity, NOVA-514151) were used during the adapter ligation step. Final library concentrations were quantified by Qubit 3 Fluorometer (Invitrogen), and library size was assessed using an Agilent 4200 TapeStation. Samples were submitted for 150 bp paired-end sequencing on a NovaSeqX at QB3 Genomics, UC Berkeley, Berkeley, CA, RRID:SCR_022170.

### mRNA-seq data analysis

Reads were aligned to the SK1 genome using HISAT2^88^ and sorted and indexed using SAMtools^89^. Transcript per million (TPM) quantifications were generated using StringTie^90^, and raw counts were generated using featureCounts^91^ for use with DESeq2.

To generate heatmaps (Fig. 2), read counts were normalized and variance stabilized using the vst() function in DESeq2 (version 1.48.2)^92^. Genes with fewer than 100 total reads across all samples were excluded prior to normalization. Variance-stabilized expression values were scaled (z-score) across all samples to visualize relative expression trends independent of absolute expression level. Genes were hierarchically clustered using Pearson correlation distance with Ward’s method (ward.D2). Heatmaps were generated using the ComplexHeatmap package (version 2.24.1) in R (version 4.5.1). Samples are grouped by condition (control, depletion) and ordered by replicate and timepoint. A “divergence” annotation was computed as the mean difference in z-scored expression (control – depletion) across the indicated timepoints, highlighting genes with TF-dependent expression during this interval.

To generate scatter plots (Fig. S2, S6A), TPM values were averaged across the two replicates within each strain and timepoint. Agreement between strains was visualized as scatter plots of mean log_2_(TPM+1) values. Correlation was quantified per timepoint as the Pearson correlation coefficient of log_2_-transformed mean TPM values. Analyses and plots were generated in R using ggplot2^93^.

For differential expression analyses comparing control and depletion conditions (Fig. 3A, C, E; 5B), relevant timepoints were first selected for the analysis based on when protein levels (Fig. 1C-E) and activity (Fig. 2, S2) of each TF peak. The raw gene counts for these timepoints were then fed into DESeq2 using the following options. Time, replicate, and condition were used as factors, with the +*OsTIR1^F74G^*condition set as the reference so genes more highly expressed in the control (-*OsTIR1^F74G^*) could be identified. The default Wald test and Benjamini-Hochberg p-value adjustment methods were used and a significance cut-off of P_adj_ ≤ 0.01 and a log_2_ fold change ≥ 0.5 were set. Any genes with fewer than 100 counts across samples were filtered out of the analysis. DESeq2 differential expression analyses were visualized using ggplot2.

For differential expression analysis to identify transcripts disproportionally affected by Msn2/Msn4/Skn7-AID depletion relative to Msn2/Msn4-AID depletion, only the *+OsTIR1^F74G^* conditions were used from each experiment. Timepoints used in this analysis (5-6 hr in SPO) were selected based on the time depletion experiments in Fig. 4F. DESeq2 was run similarly to above; here, however, the Msn2/Msn4/Skn7-AID depletion condition was set as the reference with a P_adj_ < 0.01 cut-off and no log_2_ fold change threshold. Only genes identified as being Msn2/Msn4/Skn7-dependent (Fig. 5B) were considered, though dispersion estimates and size factors were calculated using the full gene set to ensure robust empirical Bayes shrinkage of per-gene dispersion estimates. Candidate genes were further filtered based on having a larger DESeq2-caluclated log_2_ fold change in the Msn2/Msn4/Skn7-AID RNA-seq experiment (at 5-6 hr in SPO; Table S7) vs. the Msn2/Msn4-AID (at 5-6 hr in SPO; Table S6) or Skn7-AID (at 4-6 hr in SPO; Table S8) experiments. In summary, candidates identified met three conditions: 1) Msn2/Msn4/Skn7-dependent, 2) lower expression in the Msn2/Msn4/Skn7-AID depletion condition compared to the Msn2/Msn4-AID depletion condition, and 3) larger log_2_ fold change (control/depletion) in the Msn2/Msn4/Skn7-AID RNA-seq experiment than both Msn2/Msn4-AID and Skn7-AID mRNA-seq experiments during mid-gametogenesis.

GO enrichment analyses (biological process) on TF-dependent genes (Fig. S3) were performed using clusterProfiler v4.16^94,95^ with a P_adj_ < 0.05 cut-off. Redundant terms were collapsed via semantic similarity (cutoff = 0.5) using GOSemSim^96^. The chord diagram (Fig. 5E) linking *TSA1* and *AHP1* to their shared and unique GO terms (biological process, molecular function) was generated using GOChord^97^ and was limited to experimentally supported annotations retrieved from the *Saccharomyces cerevisiae* genome database.

### Sporulation efficiency, spore fitness, and germination experiments

For all gametogenesis experiments, cells were first induced to sporulate using the conditions described above and TF depletion was induced using 50 µM CuSO4 and 100 µM 5-Ph-IAA. Unless otherwise noted, depletion was induced at 2 hr in SPO. After 24 hr in SPO, sporulation efficiency was assessed using a light microscope. Portions of progenitors forming 0-1 (unsporulated/monads), 2 (dyads), or 3-4 (tri/tetrads) spores were scored (≥ 300 per experiment). Sporulation efficiency is plotted as % tri/tetrads.

Spore fitness was assessed by harvesting cells from a 24-hr SPO culture, washing twice with MilliQ H_2_O to remove CuSO_4_ and 5-Ph-IAA, digesting the asci using 1 mg/mL zymolyase (MP Biomedicals, 083209-CF) for 5 minutes at RT, then dissecting tetrads to separate individual spores on YPD plates (1%yeast extract, 2% peptone, 2% glucose, 2% agar). After 48 hr of growth at 30 °C, plates were imaged and spore colony size was quantified in FIJI by image thresholding and particle size analysis. Impaired spores were defined as colonies that were either not visible (indicating failure to germinate, or so slow growing that no colony is visible at 48 hr), or smaller than half the size of the average colony on the corresponding control (-*OsTIR1^F74G^*) plate. For each data point, 24 tetrads were dissected to assay the percentage of impaired spores out of 96 total spores. For each genotype, a minimum of two diploid strains were generated and tested.

Germination kinetics were assessed by timelapse microscopy. Sporulated cells from a 24-hr SPO culture were washed twice with MilliQ H_2_O to remove CuSO_4_ and 5-Ph-IAA, then loaded onto a Concanavalin A-treated 96-well plate (Corning, 4580). 100 µL of YPD was added and the time of emergence of the first bud was scored relative to rich media addition. Cells were imaged by wide-field microscopy on a DeltaVision Elite microscope (GE Healthcare) running the accompanying softWoRx software (version 6.5.2), using a 60x/1.42 NA oil-immersion objective and a PCO Edge sCMOS camera. Images were taken every 5 minutes for 10 hr. Acquisition was performed in a temperature-controlled chamber at 30 °C.

For heat shock gametogenesis experiments, cells were first induced to sporulate using the normal conditions described above except for that SPO cultures were started at room temperature. Depletion was induced at 1 hr to ensure full Hsf1 depletion prior to heat shock, then at 2 hr cultures were shifted to a water bath with the indicated temperature (30 °C -36 °C) and allowed to shake until 24 hr. For spore fitness, plates were always incubated at 30 °C.

Data visualization and statistical analyses were conducted using GraphPad Prism (version 11.1.0). For spore fitness and sporulation efficiency, statistical significance was determined by one-or two-way ANOVA (where indicated) with matched values and Tukey’s multiple comparisons correction using a 95% confidence interval. For germination kinetics analyses, statistical significance was determined by unpaired two-tailed t-test using a 99% confidence interval.

### Aged cell sorting and replicative lifespan analysis

The replicative lifespan of gametes from aged and young progenitors was determined as previously described^59^. Briefly, a starting population of cells was biotin labeled, then aged progenitors were later reisolated following an outgrowth step using an anti-biotin magnetic bead-sorting assay^11,98^. For each strain, an overnight YPD culture was back diluted and grown to log-phase at 30 °C. Then, 8 OD_600_ units of cells were washed three times with PBS pH 8 and labeled with 8 mg/ml EZ-Link Sulfo-NHS-LC-biotin (ThermoFisher Scientific, 21335) in PBS for 30 min at 4 °C. Cells were washed once in PBS pH 8 100 mM glycine to remove excess biotin. Biotinylated cells were grown for 16 hr in YPD with 100 μg/mL ampicillin at 30 °C, from a starting OD_600_ of 0.1. Cells were subsequently harvested, washed in PBS pH 7.4 0.5% BSA buffer, and mixed with 100 μL of anti-biotin magnetic beads (Miltenyi Biotec, 130-097-046) for 15 min at 4 °C. Cells were washed with PBS pH 7.4 0.5% BSA buffer and sorted magnetically using LS depletion columns with a QuadroMacs sorter, washing three times with PBS pH 7.4 0.5% BSA. This magnetic sorting was then repeated a second time to increase purity of aged cells. A fraction of the initial flow-through (biotin-negative) was kept as young cells alongside eluted (biotin-positive) aged cells.

Both young and aged cells were labeled for 15 min at RT using 1 μg/mL Wheat Germ Agglutinin, Alexa Fluor 350 Conjugate (ThermoFisher Scientific, W11263) to visualize bud scars, and 1 µg/mL anti-streptavidin 564 nm dye to visualize biotinylation. 0.1 OD_600_ units of cells were fixed in 3.7% formaldehyde to and imaged to assess pre-sporulation age by counting biotinylated cells and bud scars. The remaining young and aged cells were subsequently washed three times with SPO media, then resuspended in conditioned SPO at a cell density of OD_600_ = 0.8 with 100 μg/mL ampicillin and incubated for 24 hr with shaking at 30 °C in the dark. At 2 hr in SPO, TF depletion was induced using 50 µM CuSO_4_ and 100 µM 5-Ph-IAA. Again, 0.1 OD_600_ units of cells were fixed after the 24-hr SPO culture to assess post-sporulation age by counting biotinylated cells and bud scars.

Post-sporulation age-sorted cultures were pelleted and resuspended in 1 mL digestion buffer (0.1 mg/mL zymolyase and 2% β-mercaptoethanol in H_2_O), then incubated for 3 hr on a nutator at 30 °C. Cells were washed twice and resuspended in H_2_O, then separated by repeated sonication (6x 20 seconds, 50% pulses) at 4 °C. Upon separation, spores were filtered (30 µm pre-separation filter, Militenyi Biotec, 130-041-407) and diluted in YPD + ampicillin (100 μg/mL) to an OD_600_ of 0.8. After 3 hr outgrowth at 30 °C to increase gamete cell size, 1 mL of cell suspension was aliquoted, sonicated, filtered, and loaded into the microfluidics system.

Microfluidics devices (adapted from^99^; 18++ chips from iBiochips) were set up using the manufacturer’s recommendations on an ECHO Revolution fluorescence microscope in the inverted mode with the environmental chamber set to 30 °C. Briefly, 20 mL syringes (Becton, Dickinson and Company) filled with ∼15 mL of YPD 2% + ampicillin (100 μg/mL) + 1% Pluronic F-127 surfactant (P2443, Sigma) were connected to a Clay Adams Intramedic Luer-Stub Adapter (Becton, Dickinson and Company). Non-DEHP Medical Grade Tubing (iBiochips, MTB-100, ID = 0.202”, OD = 0.060”, wall = 0.20”) equipped with a hollow stainless-steel pin (iBiochips, HSP-200) was then used to connect the media syringe to the imaging units using the “media loading” ports. “Cell loading” ports were plugged with solid stainless-steel pins (iBiochips, SSP-200) while imaging units were primed with media for at least 2 hr with a flow rate of 2.4 μL/min using micropumps (iBiochips,78-7200BIO). After all units were filled with media, pumps were stopped an hour prior to cell loading. For cell loading, 5 mL syringes (Becton, Dickinson and Company) with Clay Adams Intramedic Luer-Stub Adapter (Becton, Dickinson and Company) were filled with cells diluted to an OD_600_ of 0.8 in YPD + ampicillin (100 μg/mL). Solid stainless-steel pins were then removed from the “cell loading” ports and tubing and open stainless-steel pins were used to connect the syringe containing cells with the microfluidic device. Cells were loaded manually and, once sufficient cells were loaded, the “cell loading” ports were plugged with solid stainless-steel pins and media flow was resumed at 2.4 μL/min. Timelapse imaging was performed using an Olympus 40x/1.4 NA oil-immersion objective, at 15–20-minute intervals for 72 hr. Image acquisition of chamber was achieved with 2×27 tiled fields of view, with three 3-5 µm z-slices.

After a movie was completed and processed, single Pma2-GFP-positive cells (gametes) were identified and cropped. Next, the replicative lifespan of individual gametes was assessed in the brightfield channel by counting the number of daughter cells produced.

---

**Fig. S1.**
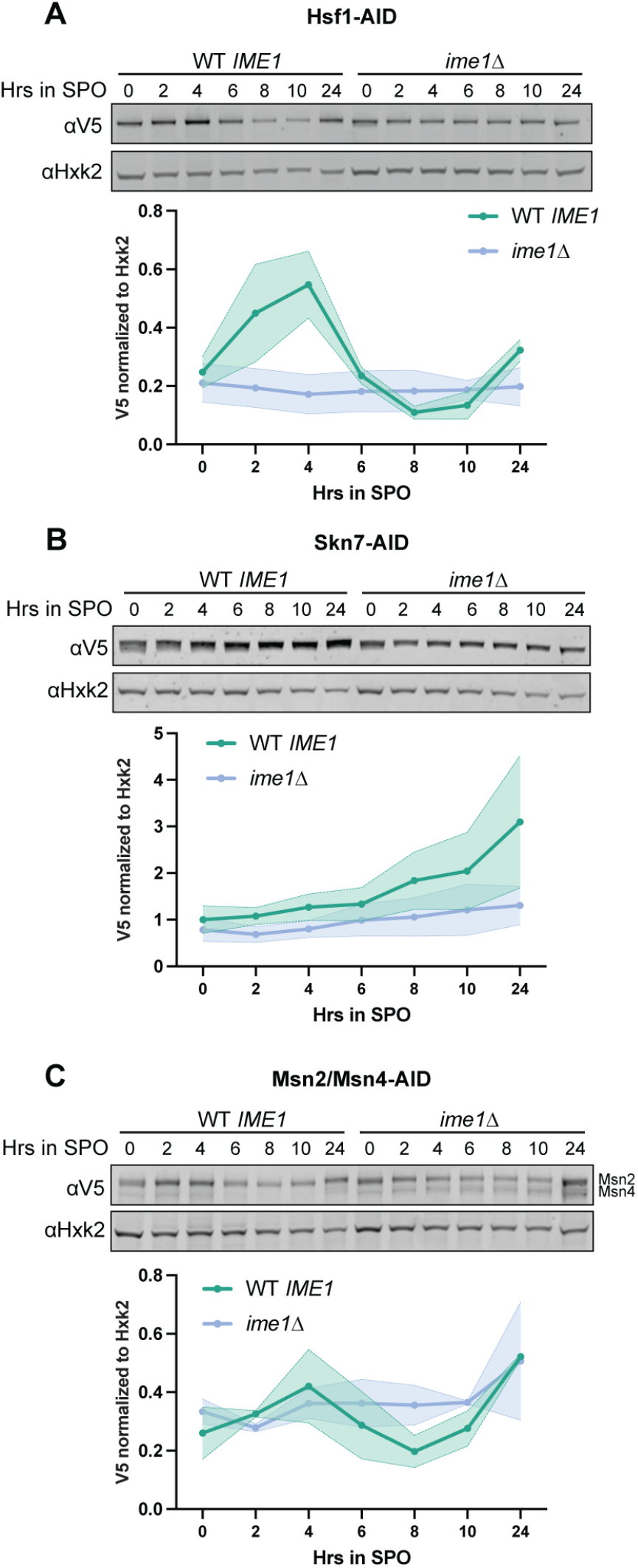
Stress-responsive TFs exhibit dynamic, gametogenesis-specific expression. **A-C** Western blots showing Hsf1-AID (A), Skn7-AID (B), and Msn2/Msn4-AID (C) protein levels throughout gametogenesis in WT *IME1* and *ime1Δ* backgrounds. Hxk2 serves as a loading control. Quantification below blot, shown is the average of two replicates, with shaded region representing the range.

**Fig. S2.**
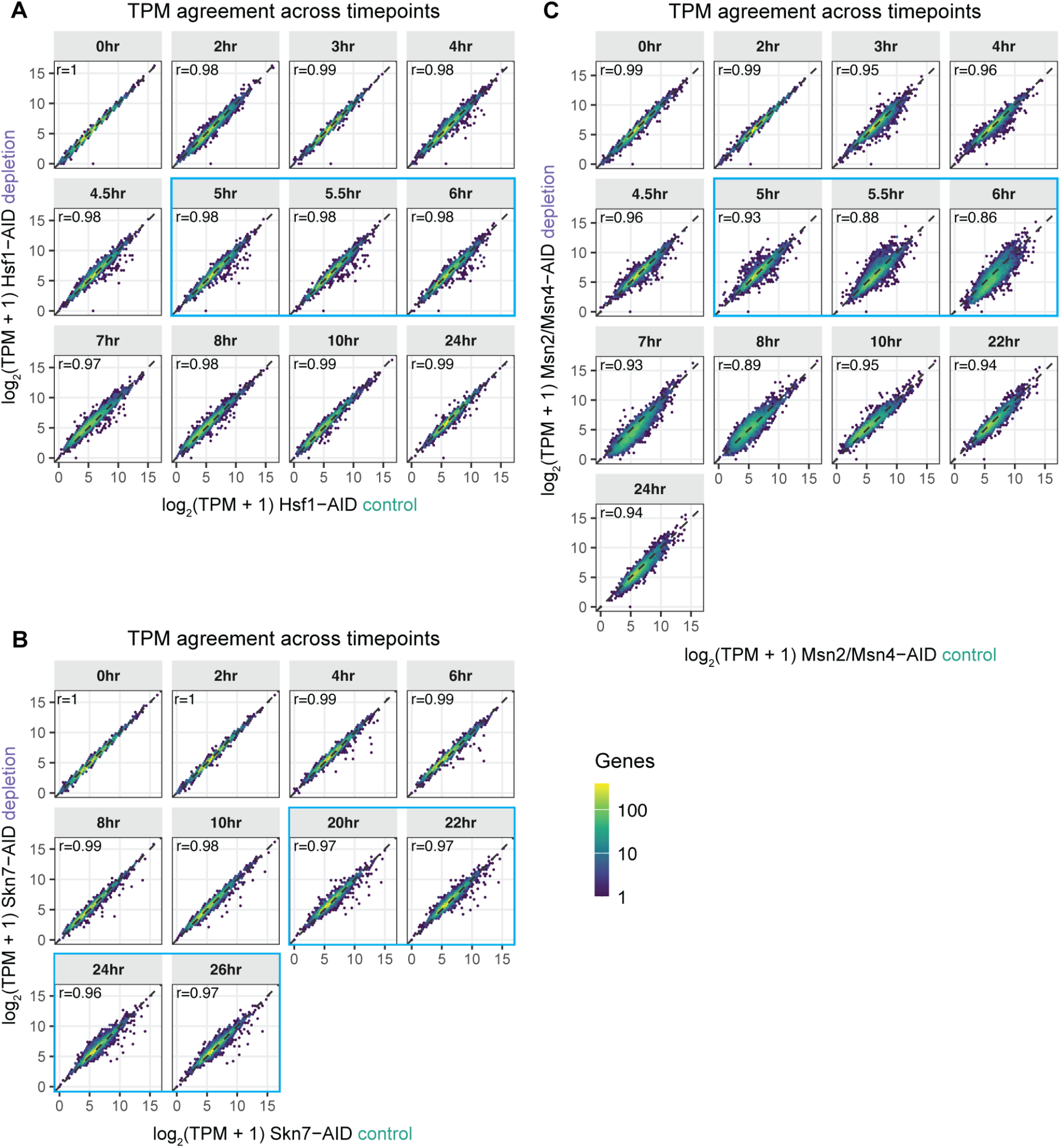
TF depletion does not cause widespread global gene expression changes. **A-C** Scatter plots comparing log_2_-transformed transcript abundance (TPM +1) for each gene in control (x-axis) versus depletion (y-axis) conditions at each sampled timepoint (hr in SPO) for Hsf1-AID (A), Skn7-AID (B), and Msn2/Msn4-AID (C). TPM values are averaged across two replicates. Hexagons represent bins of genes and are colored by the number of genes they contain on a log_10_ scale (dark to light indicating fewer to more genes). The dashed line indicates y = x (perfect agreement). Hexagons below the dashed line indicate genes with lower expression upon TF depletion, consistent with TF-dependent expression. Pearson correlation coefficients (*r*) calculated on log_2_(TPM + 1) values for all genes are shown in the upper left of each panel. Blue boxes indicate timepoints used to identify differentially expressed (TF-dependent) genes with DESeq2.

**Fig. S3.**
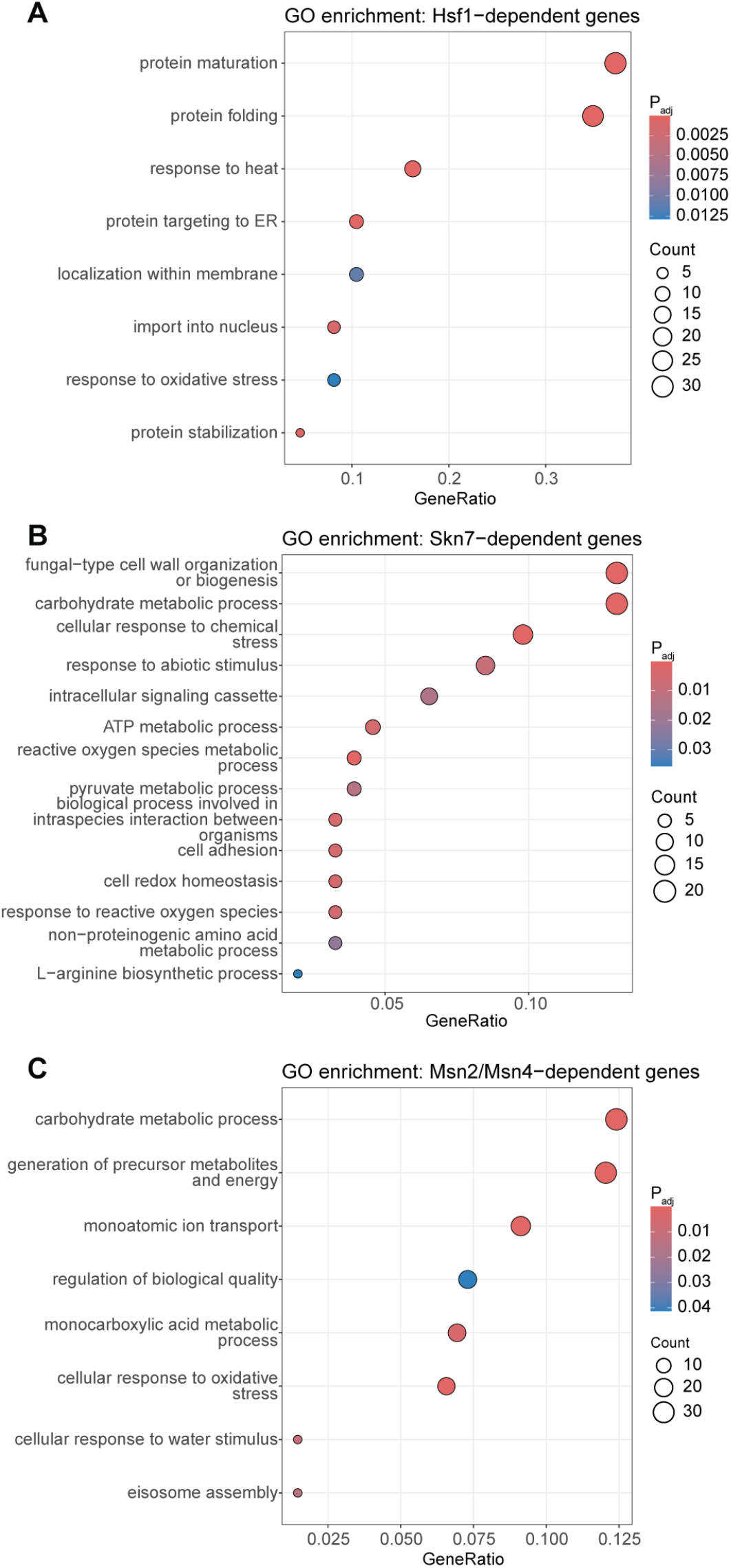
TF-dependent genes during gametogenesis are enriched for stress response, quality control, and homeostasis genes. **A-C** GO enrichment dot plots for meiotic Hsf1-(A), Skn7-(B), and Msn2/Msn4-(C) dependent genes. Top categories with P_adj_ < 0.05 are shown with redundant terms collapsed (see Methods).

**Fig. S4.**
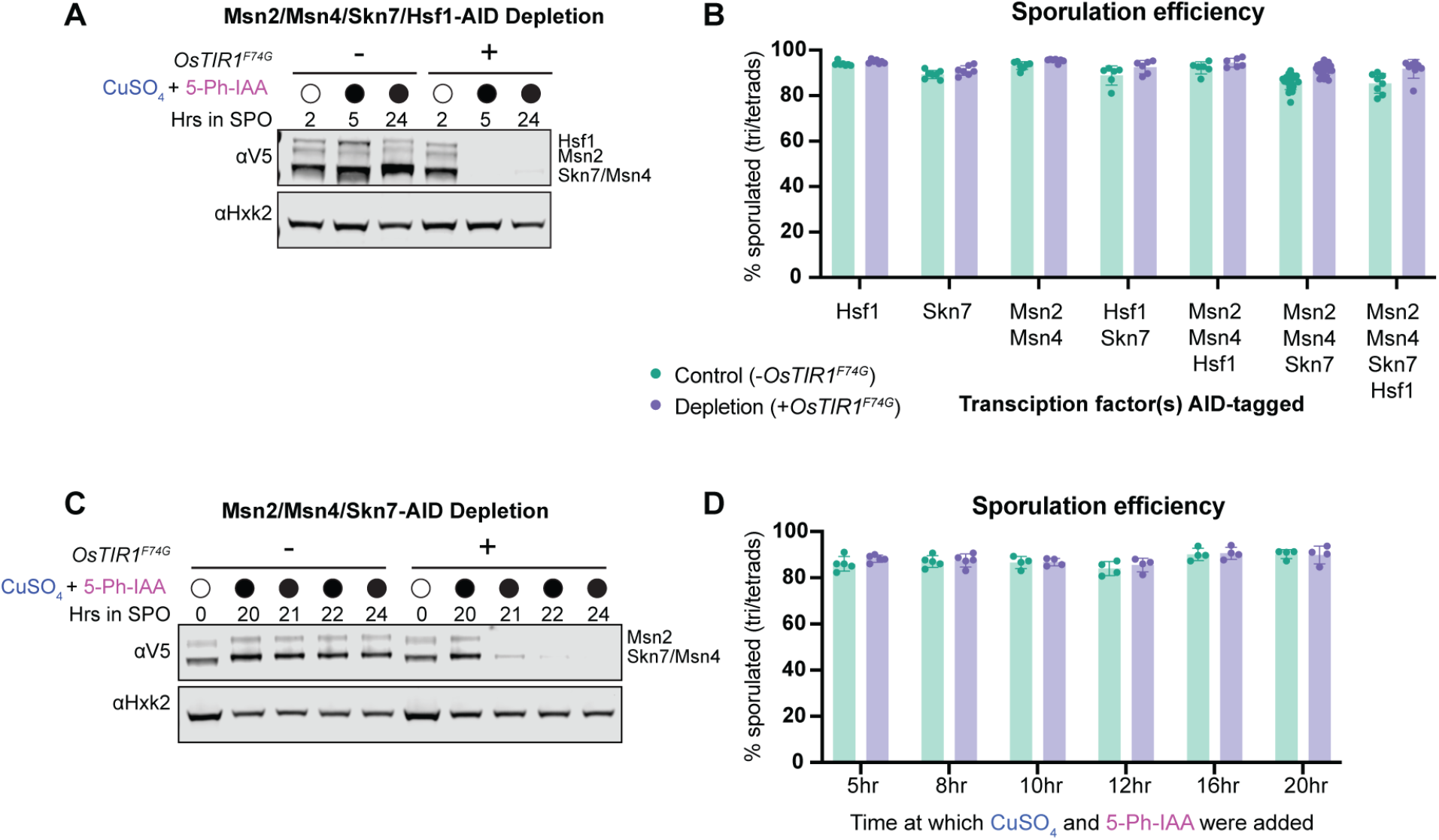
Multiple stress-responsive TFs are robustly depleted simultaneously without reducing sporulation efficiency. **A** Western blot showing Msn2/Msn4/Skn7/Hsf1-AID protein levels during gametogenesis in control (-*OsTIR1^F74G^*) and depletion (+*OsTIR1^F74G^*) conditions. Msn4-AID and Skn7-AID are only 1 kDa apart in size and are indistinguishable on the blot. Depletion was induced at 2 hr in SPO by addition of 50 µM CuSO_4_ and 100 µM 5-Ph-IAA. Hxk2 serves as a loading control. **B** Sporulation efficiency results from individual and combinatorial TF-AID depletion experiments in Fig. 4B. Shown is the average ± standard deviation (n ≥ 6). **C** Western blot showing Msn2/Msn4/Skn7-AID protein levels during gametogenesis in control (-*OsTIR1^F74G^*) and depletion (+*OsTIR1^F74G^*) conditions. Msn4-AID and Skn7-AID are only 1 kDa apart in size and are indistinguishable on the blot. Depletion was induced at 20 hr in SPO by addition of 50 µM CuSO_4_ and 100 µM 5-Ph-IAA. Hxk2 serves as a loading control. **D** Sporulation efficiency results from Msn2/Msn4/Skn7-AID time-varied depletion experiments in Fig. 4F. Shown is the average ± standard deviation (n ≥ 4).

**Fig. S5.**
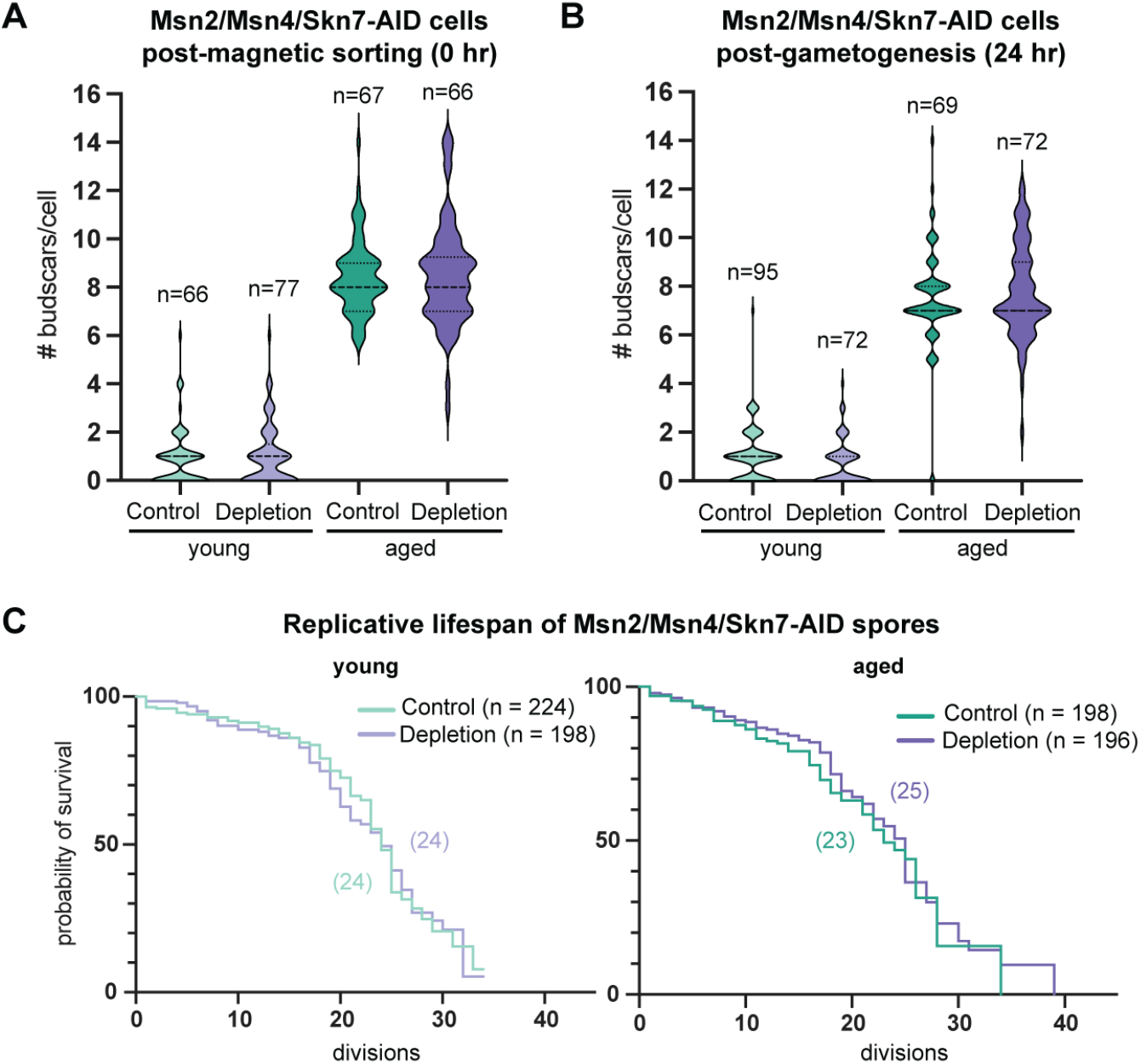
Msn2, Msn4, and Skn7 depletion during gametogenesis does not affect spore replicative lifespan. **A-B** Quantification of number of budscars/cell as a proxy for age in cells directly after aging and sorting (A) and after going through gametogenesis (B). **C** Replicative lifespan of spores from young (left) and aged (right) Msn2/Msn4/Skn7-AID progenitor cells under control and depletion conditions. Median lifespan shown in parentheses next to each curve.

**Fig. S6.**
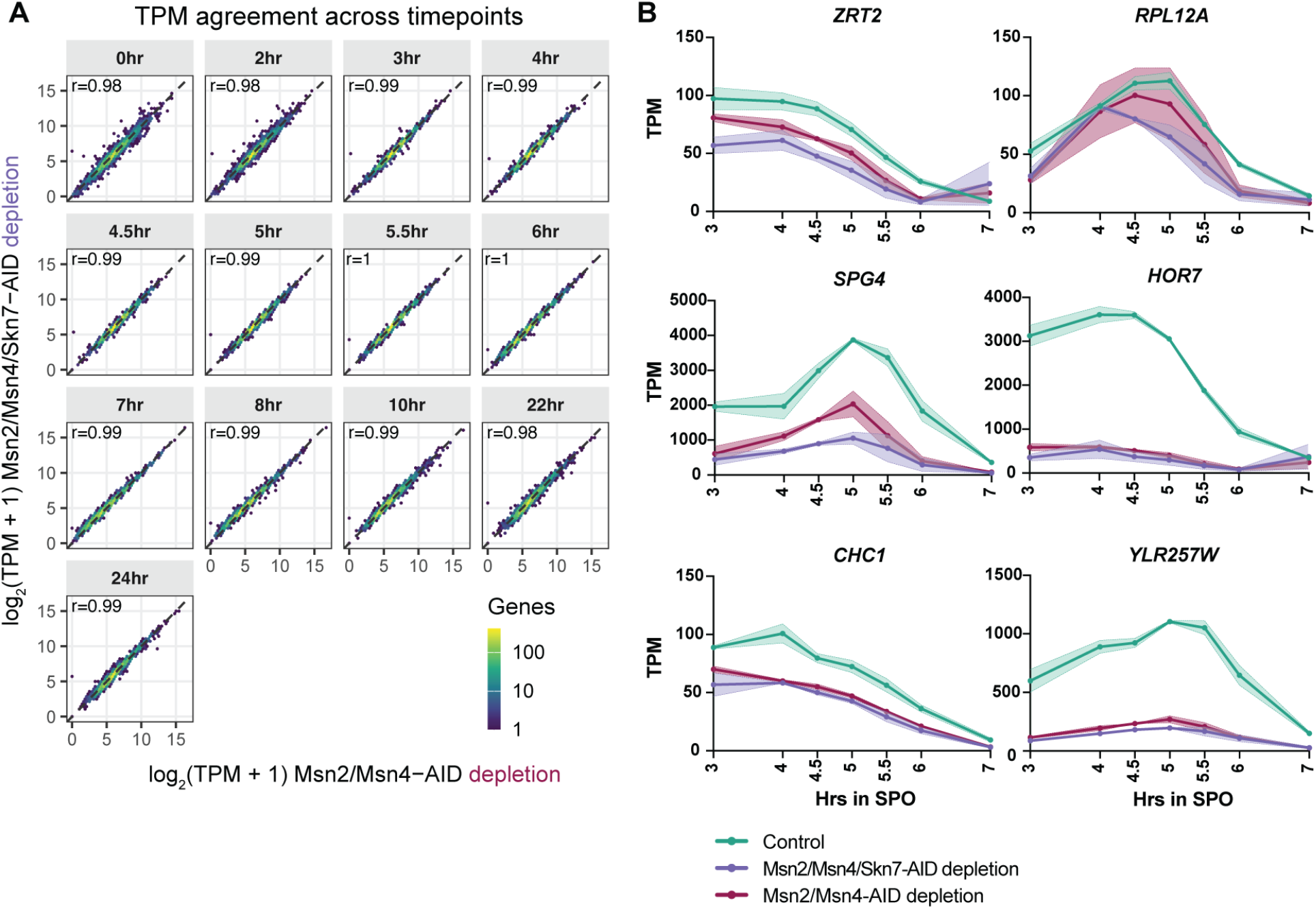
Transcriptional differences between Msn2/Msn4-AID and Msn2/Msn4/Skn7-AID depletion conditions. **A** Scatter plots comparing log_2_-transformed transcript abundance (TPM +1) for each gene in Msn2/Msn4-AID depletion (x-axis) versus Msn2/Msn4/Skn7-AID depletion (y-axis) conditions, with each panel corresponding to a different meiotic timepoint (hr in SPO). TPM values are averaged across two replicates. Hexagons represent bins of genes and are colored by the number of genes they contain on a log_10_ scale (dark to light indicating fewer to more genes). The dashed line indicates y = x (perfect agreement). Pearson correlation coefficients (*r*) calculated on log_2_(TPM + 1) values for all genes are shown in the upper left of each panel. **B** Plots showing TPM levels during gametogenesis of genes (other than *TSA1* and *AHP1*) identified by analysis in Fig. 5C. Shown is the average of two replicates, with shaded region representing the range. Control is from the Msn2/Msn4/Skn7-AID experiment.

**Fig. S7.**
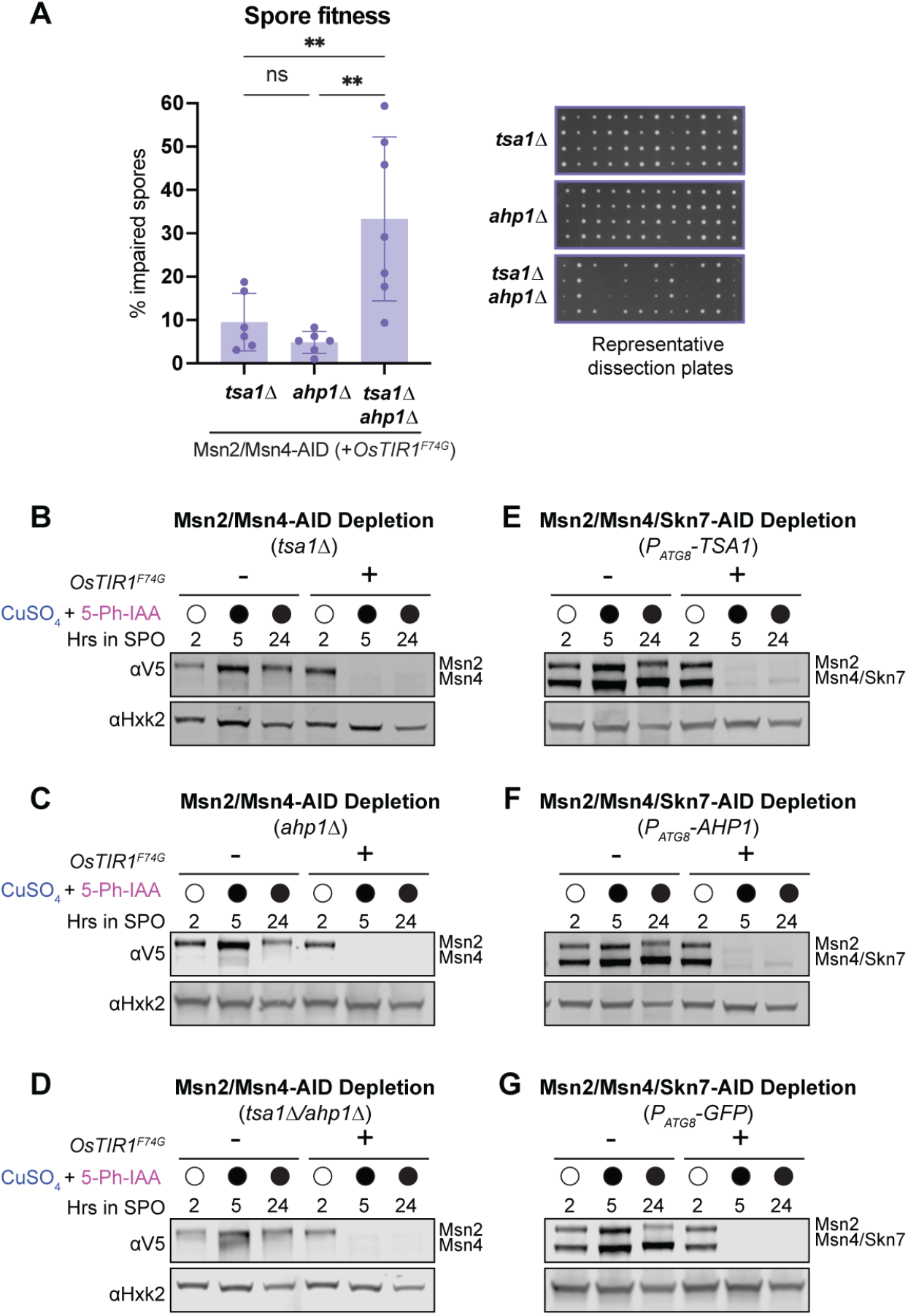
*TSA1* and *AHP1* promote gamete fitness. **A** Results from spore fitness experiments with deletion strains in Msn2/Msn4-AID depletion background. Experiment conducted as in Fig. 4A. Shown is the average ± standard deviation (n ≥ 6). Statistical significance determined by one-way ANOVA with Tukey’s multiple comparisons correction (** P_adj_ < 0.01). Representative dissection plates from each genotype shown on right. **B-D** Western blots showing Msn2/Msn4-AID protein levels during gametogenesis in control (-*OsTIR1^F74G^*) and depletion (+*OsTIR1^F74G^*) conditions in *tsa1Δ* (B), *ahp1Δ* (C), and *tsa1Δ ahp1Δ* (D) strains. Depletion was induced at 2 hr in SPO by addition of 50 µM CuSO_4_ and 100 µM 5-Ph-IAA. Hxk2 serves as a loading control. **E-G** Western blots showing Msn2/Msn4/Skn7-AID protein levels during gametogenesis in control (-*OsTIR1^F74G^*) and depletion (+*OsTIR1^F74G^*) conditions in *P_ATG8_-TSA1* (E), *P_ATG8_-AHP1* (F), and *P_ATG8_-GFP* (G) strains. Depletion was induced at 2 hr in SPO by addition of 50 µM CuSO_4_ and 100 µM 5-Ph-IAA. Hxk2 serves as a loading control.

## References

1. Marston, A. L. & Amon, A. Meiosis: cell-cycle controls shuffle and deal. Nat. Rev. Mol. Cell Biol. 5, 983–997 (2004).

2. Goodman, J. S., King, G. A. & Ünal, E. Cellular quality control during gametogenesis. Exp. Cell Res. 396, 112247 (2020).

3. Xiao, T. & Ünal, E. Remodeling, compartmentalization, and degradation: a trifecta for organelle quality control during gametogenesis. Curr. Opin. Genet. Dev. 92, 102347 (2025).

4. Chu, S. et al. The Transcriptional Program of Sporulation in Budding Yeast. Science 282, 699–705 (1998).

5. Neiman, A. M. Sporulation in the Budding Yeast *Saccharomyces cerevisiae*. Genetics 189, 737–765 (2011).

6. Brar, G. A. et al. High-Resolution View of the Yeast Meiotic Program Revealed by Ribosome Profiling. Science 335, 552–557 (2012).

7. Suda, Y. et al. Remodeling of the secretory pathway is coordinated with de novo membrane formation in budding yeast gametogenesis. iScience 27, 110855 (2024).

8. Suda, Y., Nakanishi, H., Mathieson, E. M. & Neiman, A. M. Alternative Modes of Organellar Segregation during Sporulation in *Saccharomyces cerevisiae*. Eukaryot. Cell 6, 2009–2017 (2007).

9. Sawyer, E. M. et al. Developmental regulation of an organelle tether coordinates mitochondrial remodeling in meiosis. J. Cell Biol. 218, 559–579 (2019).

10. Otto, G. M., Cheunkarndee, T., Leslie, J. M. & Brar, G. A. Programmed cortical ER collapse drives selective ER degradation and inheritance in yeast meiosis. J. Cell Biol. 220, e202108105 (2021).

11. King, G. A. et al. Meiotic cellular rejuvenation is coupled to nuclear remodeling in budding yeast. eLife 8, e47156 (2019).

12. Hsu, T.-H., Chen, R.-H., Cheng, Y.-H. & Wang, C.-W. Lipid droplets are central organelles for meiosis II progression during yeast sporulation. Mol. Biol. Cell 28, 440– 451 (2017).

13. Sing, T. L., Brar, G. A. & Ünal, E. Gametogenesis: Exploring an Endogenous Rejuvenation Program to Understand Cellular Aging and Quality Control. Annu. Rev. Genet. 56, 89– 112 (2022).

14. Jambhekar, A. & Amon, A. Control of Meiosis by Respiration. Curr. Biol. 18, 969–975 (2008).

15. Causton, H. C. et al. Remodeling of Yeast Genome Expression in Response to Environmental Changes. Mol. Biol. Cell 12, 323–337 (2001).

16. Gasch, A. P. et al. Genomic Expression Programs in the Response of Yeast Cells to Environmental Changes□D. Mol. Biol. Cell 11, (2000).

17. Gasch, A. P. Twenty-Five Years of the Environmental Stress Response and the Enduring Power of Yeast in Stress Biology. Yeast 43, 189–201 (2026).

18. Hu, Z., Killion, P. J. & Iyer, V. R. Genetic reconstruction of a functional transcriptional regulatory network. Nat. Genet. 39, 683–687 (2007).

19. Morano, K. A., Grant, C. M. & Moye-Rowley, W. S. The Response to Heat Shock and Oxidative Stress in *Saccharomyces cerevisiae*. Genetics 190, 1157–1195 (2012).

20. Verghese, J., Abrams, J., Wang, Y. & Morano, K. A. Biology of the Heat Shock Response and Protein Chaperones: Budding Yeast (Saccharomyces cerevisiae) as a Model System. Microbiol. Mol. Biol. Rev. 76, 115–158 (2012).

21. Li, J., Chauve, L., Phelps, G., Brielmann, R. M. & Morimoto, R. I. E2F coregulates an essential HSF developmental program that is distinct from the heat-shock response. Genes Dev. 30, 2062–2075 (2016).

22. Reimold, A. M. et al. Plasma cell differentiation requires the transcription factor XBP-1. Nature 412, 300–307 (2001).

23. Gass, J. N., Gifford, N. M. & Brewer, J. W. Activation of an Unfolded Protein Response during Differentiation of Antibody-secreting B Cells. J. Biol. Chem. 277, 49047–49054 (2002).

24. Cheng, Z. et al. Pervasive, Coordinated Protein-Level Changes Driven by Transcript Isoform Switching during Meiosis. Cell 172, 910–923.e16 (2018).

25. Bartolutti, C. et al. The UPR drives gametogenesis via an unexpected response, coordinating translation and ER structure. J. Cell Biol. 225, e202509100 (2026).

26. Yesbolatova, A. et al. The auxin-inducible degron 2 technology provides sharp degradation control in yeast, mammalian cells, and mice. Nat. Commun. 11, 5701 (2020).

27. Van Werven, F. J. & Amon, A. Regulation of entry into gametogenesis. Philos. Trans. R. Soc. B Biol. Sci. 366, 3521–3531 (2011).

28. Chia, M. & Van Werven, F. J. Temporal Expression of a Master Regulator Drives Synchronous Sporulation in Budding Yeast. G3 GenesGenomesGenetics 6, 3553–3560 (2016).

29. Boy-Marcotte, E. et al. The heat shock response in yeast: differential regulations and contributions of the Msn2p/Msn4p and Hsf1p regulons. Mol. Microbiol. 33, 274–283 (1999).

30. Pincus, D. et al. Genetic and epigenetic determinants establish a continuum of Hsf1 occupancy and activity across the yeast genome. Mol. Biol. Cell 29, 3168–3182 (2018).

31. Ciccarelli, M., Masser, A. E., Kaimal, J. M., Planells, J. & Andréasson, C. Genetic inactivation of essential *HSF1* reveals an isolated transcriptional stress response selectively induced by protein misfolding. Mol. Biol. Cell 34, ar101 (2023).

32. Solís, E. J. et al. Defining the Essential Function of Yeast Hsf1 Reveals a Compact Transcriptional Program for Maintaining Eukaryotic Proteostasis. Mol. Cell 63, 60–71 (2016).

33. Hasan, R. et al. The control of the yeast H2O2 response by the Msn2/4 transcription factors. Mol. Microbiol. 45, 233–241 (2002).

34. Martínez-Pastor, M. T. et al. The Saccharomyces cerevisiae zinc finger proteins Msn2p and Msn4p are required for transcriptional induction through the stress response element (STRE). EMBO J. 15, 2227–2235 (1996).

35. Brombacher, K., Fischer, B. B., Rüfenacht, K. & Eggen, R. I. L. The role of Yap1p and Skn7p-mediated oxidative stress response in the defence of *Saccharomyces cerevisiae* against singlet oxygen. Yeast 23, 741–750 (2006).

36. Lee, J. et al. Yap1 and Skn7 Control Two Specialized Oxidative Stress Response Regulons in Yeast. J. Biol. Chem. 274, 16040–16046 (1999).

37. Godon, C. et al. The H2O2 Stimulon in Saccharomyces cerevisiae. J. Biol. Chem. 273, 22480–22489 (1998).

38. Jamieson, D. J. Oxidative stress responses of the yeast Saccharomyces cerevisiae. 17 (1998).

39. Morgan, B. A. The Skn7 response regulator controls gene expression in the oxidative stress response of the budding yeast Saccharomyces cerevisiae. EMBO J. 16, 1035– 1044 (1997).

40. Mühlhofer, M. et al. Deletion of the transcription factors Hsf1, Msn2 and Msn4 in yeast uncovers transcriptional reprogramming in response to proteotoxic stress. FEBS Lett. 598, 635–657 (2024).

41. Werner-Washburne, M., Stone, D. E. & Craig, E. A. Complex Interactions among Members of an Essential Subfamily of hsp7O Genes in Saccharomyces cerevisiae. MOL CELL BIOL.

42. Caplan, A. J. & Douglas, M. G. Characterization of YDJI: A Yeast Homologue of the Bacterial dnaJ Protein. J. Cell Biol. 114, (1991).

43. Luke, M. M., Sutton, A. & Arndt, K. T. Characterization of SISI, a Saccharomyces cerevisiae Homologue of Bacterial dnaJ Proteins. J. Cell Biol. 114, (1991).

44. Leonhardt, S. A. & Fearon, K. HSP78 Encodes a Yeast Mitochondrial Heat Shock Protein in the Clp Family of ATP-Dependent Proteases. MOL CELL BIOL.

45. Manning, B. D., Padmanabha, R. & Snyder, M. The Rho-GEF Rom2p Localizes to Sites of Polarized Cell Growth and Participates in Cytoskeletal Functions in *Saccharomyces cerevisiae*. Mol. Biol. Cell 8, 1829–1844 (1997).

46. Yin, Q. Y., et al. Comprehensive Proteomic Analysis of Saccharomyces cerevisiae Cell Walls. J. Biol. Chem. 280, 20894–20901 (2005).

47. Van Der Vaart, J. M., Caro, L. H., Chapman, J. W., Klis, F. M. & Verrips, C. T. Identification of three mannoproteins in the cell wall of Saccharomyces cerevisiae. J. Bacteriol. 177, 3104–3110 (1995).

48. Cohen, B. A., Pilpel, Y., Mitra, R. D. & Church, G. M. Discrimination between Paralogs using Microarray Analysis: Application to the Yap1p and Yap2p Transcriptional Networks. Mol. Biol. Cell 13, 1608–1614 (2002).

49. Venters, B. J. et al. A Comprehensive Genomic Binding Map of Gene and Chromatin Regulatory Proteins in Saccharomyces. Mol. Cell 41, 480–492 (2011).

50. Bossier, P., Fitch, I. T., Boucherie, H. & Tuite, M. F. Structure and expression of a yeast gene encoding the small heat-shock protein Hsp26. Gene 78, 323–330 (1989).

51. Praekelt, U. M. & Meacock, P. A. HSP12, a new small heat shock gene of Saccharomyces cerevMae: Analysis of structure, regulation and function.

52. Kim, S. X., Çamdere, G., Hu, X., Koshland, D. & Tapia, H. Synergy between the small intrinsically disordered protein Hsp12 and trehalose sustain viability after severe desiccation. eLife 7, e38337 (2018).

53. Sales, K., Brandt, W., Rumbak, E. & Lindsey, G. The LEA-like protein HSP 12 in Saccharomyces cerevisiae has a plasma membrane location and protects membranes against desiccation and ethanol-induced stress. Biochim. Biophys. Acta BBA -Biomembr. 1463, 267–278 (2000).

54. Stanford, D. R. et al. Division of Labor Among the Yeast Sol Proteins Implicated in tRNA Nuclear Export and Carbohydrate Metabolism. Genetics 168, 117–127 (2004).

55. Rintala, E., Pitkänen, J., Vehkomäki, M., Penttilä, M. & Ruohonen, L. The ORF *YNL274c* ( *GOR1* ) codes for glyoxylate reductase in *Saccharomyces cerevisiae*. Yeast 24, 129–136 (2007).

56. Hwang, P. K., Tugendreich, S. & Fletterick, R. J. Molecular Analysis of GPHI, the Gene Encoding Glycogen Phosphorylase in Saccharomyces cerevisiae. 9, (1989).

57. Hugener, J. et al. FilamentID reveals the composition and function of metabolic enzyme polymers during gametogenesis. Cell 187, 3303–3318.e18 (2024).

58. Unal, E., Kinde, B. & Amon, A. Gametogenesis Eliminates Age-Induced Cellular Damage and Resets Life Span in Yeast. Science 332, 1554–1557 (2011).

59. Spiri, S., et al. A role for selective autophagy of the ER in gametogenic rejuvenation revealed by microfluidics-based lifespan profiling. (2025).

60. Sorger, P. K. & Pelham, H. R. B. Yeast heat shock factor is an essential DNA-binding protein that exhibits temperature-dependent phosphorylation. Cell 54, 855–864 (1988).

61. Abane, R. & Mezger, V. Roles of heat shock factors in gametogenesis and development. FEBS J. 277, 4150–4172 (2010).

62. The Role of Heat Shock Proteins in Reproductive System Development and Function. vol. 222 (Springer International Publishing, Cham, 2017).

63. Bomblies, K., Higgins, J. D. & Yant, L. Meiosis evolves: adaptation to external and internal environments. New Phytol. 208, 306–323 (2015).

64. Morgan, C. H., Zhang, H. & Bomblies, K. Are the effects of elevated temperature on meiotic recombination and thermotolerance linked via the axis and synaptonemal complex? Philos. Trans. R. Soc. B Biol. Sci. 372, 20160470 (2017).

65. Zhang, R. et al. Rim4-seeded stress granules connect temperature sensing to meiotic regulation. Nat. Commun. 16, 5566 (2025).

66. De Storme, N. & Geelen, D. The impact of environmental stress on male reproductive development in plants: biological processes and molecular mechanisms. Plant Cell Environ. 37, 1–18 (2014).

67. Chae, H. Z., Kim, I. H., Kim, K. & Rhee, S. G. Cloning, sequencing, and mutation of thiol-specific antioxidant gene of Saccharomyces cerevisiae. J. Biol. Chem. 268, 16815– 16821 (1993).

68. Trotter, E. W., Rand, J. D., Vickerstaff, J. & Grant, C. M. The yeast Tsa1 peroxiredoxin is a ribosome-associated antioxidant. Biochem. J. 412, 73–80 (2008).

69. Jang, H. H. et al. Two Enzymes in One: Two Yeast Peroxiredoxins Display Oxidative Stress-Dependent Switching from a Peroxidase to a Molecular Chaperone Function. Cell 117, 625–635 (2004).

70. Zimmermann, J. et al. Tsa1 is the dominant peroxide scavenger and a source of H2O2-dependent GSSG production in yeast. Free Radic. Biol. Med. 226, 408–420 (2025).

71. Lee, J., Spector, D., Godon, C., Labarre, J. & Toledano, M. B. A New Antioxidant with Alkyl Hydroperoxide Defense Properties in Yeast. J. Biol. Chem. 274, 4537–4544 (1999).

72. Gao, J., Qin, Y. & Schimenti, J. C. Gene regulation during meiosis. Trends Genet. 40, 326–336 (2024).

73. Chu, S. & Herskowitz, I. Gametogenesis in Yeast Is Regulated by a Transcriptional Cascade Dependent on Ndt80. Mol. Cell 1, 685–696 (1998).

74. Kassir, Y., Granot, D. & Simchen, G. IME1, a positive regulator gene of meiosis in S. cerevisiae. Cell 52, 853–862 (1988).

75. Lindgren, A. The pachytene checkpoint in Saccharomyces cerevisiae requires the Sum1 transcriptional repressor. EMBO J. 19, 6489–6497 (2000).

76. Vershon, A. K. & Pierce, M. Transcriptional regulation of meiosis in yeast. Curr. Opin. Cell Biol. 12, 334–339 (2000).

77. He, X.-J., Mulford, K. E. & Fassler, J. S. Oxidative Stress Function of the *Saccharomyces cerevisiae* Skn7 Receiver Domain. Eukaryot. Cell 8, 768–778 (2009).

78. He, X. & Fassler, J. S. Identification of novel Yap1p and Skn7p binding sites involved in the oxidative stress response of *Saccharomyces cerevisiae*. Mol. Microbiol. 58, 1454– 1467 (2005).

79. Toledano, M. B., Kumar, C., Le Moan, N., Spector, D. & Tacnet, F. The system biology of thiol redox system in *Escherichia coli* and yeast: Differential functions in oxidative stress, iron metabolism and DNA synthesis. FEBS Lett. 581, 3598–3607 (2007).

80. Kim, K., Kim, I. H., Lee, K. Y., Rhee, S. G. & Stadtman, E. R. The isolation and purification of a specific “protector” protein which inhibits enzyme inactivation by a thiol/Fe(III)/O2 mixed-function oxidation system. J. Biol. Chem. 263, 4704–4711 (1988).

81. Aitken, R. J. Impact of oxidative stress on male and female germ cells: implications for fertility. Reproduction 159, R189–R201 (2020).

82. Bähler, J. et al. Heterologous modules for efficient and versatile PCR-based gene targeting inSchizosaccharomyces pombe. Yeast 14, 943–951 (1998).

83. Knop, M. et al. Epitope tagging of yeast genes using a PCR-based strategy: more tags and improved practical routines. Yeast 15, 963–972 (1999).

84. Daniel Gietz, R. & Woods, R. A. Transformation of yeast by lithium acetate/single-stranded carrier DNA/polyethylene glycol method. in Methods in Enzymology vol. 350 87–96 (Elsevier, 2002).

85. King, G. A. et al. Meiotic nuclear pore complex remodeling provides key insights into nuclear basket organization. J. Cell Biol. 222, e202204039 (2023).

86. Harris, A. The transcriptional regulator Ume6 is a major driver of early gene expression during gametogenesis.

87. Nishimura, K., Fukagawa, T., Takisawa, H., Kakimoto, T. & Kanemaki, M. An auxin-based degron system for the rapid depletion of proteins in nonplant cells. Nat. Methods 6, 917–922 (2009).

88. Kim, D., Paggi, J. M., Park, C., Bennett, C. & Salzberg, S. L. Graph-based genome alignment and genotyping with HISAT2 and HISAT-genotype. Nat. Biotechnol. 37, 907– 915 (2019).

89. Li, H. et al. The Sequence Alignment/Map format and SAMtools. Bioinformatics 25, 2078–2079 (2009).

90. Pertea, M. et al. StringTie enables improved reconstruction of a transcriptome from RNA-seq reads. Nat. Biotechnol. 33, 290–295 (2015).

91. Liao, Y., Smyth, G. K. & Shi, W. featureCounts: an efficient general purpose program for assigning sequence reads to genomic features. Bioinformatics 30, 923–930 (2014).

92. Love, M. I., Huber, W. & Anders, S. Moderated estimation of fold change and dispersion for RNA-seq data with DESeq2. Genome Biol. 15, 550 (2014).

93. Wickham, H. Ggplot2: Elegant Graphics for Data Analysis. (Springer-Verlag New York, 2016).

94. Xu, S. et al. Using clusterProfiler to characterize multiomics data. Nat. Protoc. 19, 3292–3320 (2024).

95. Yu, G., Wang, L.-G., Han, Y. & He, Q.-Y. clusterProfiler: an R Package for Comparing Biological Themes Among Gene Clusters. OMICS J. Integr. Biol. 16, 284–287 (2012).

96. Yu, G. et al. GOSemSim: an R package for measuring semantic similarity among GO terms and gene products. Bioinformatics 26, 976–978 (2010).

97. Walter, W., Sánchez-Cabo, F. & Ricote, M. GOplot: an R package for visually combining expression data with functional analysis. Bioinformatics 31, 2912–2914 (2015).

98. Smeal, T., Claus, J., Kennedy, B., Cole, F. & Guarente, L. Loss of Transcriptional Silencing Causes Sterility in Old Mother Cells of S. cerevisiae. Cell 84, 633–642 (1996).

99. Jo, M. C., Liu, W., Gu, L., Dang, W. & Qin, L. High-throughput analysis of yeast replicative aging using a microfluidic system. Proc. Natl. Acad. Sci. 112, 9364–9369 (2015).

